# Flytrap-Inspired Mesh-Trap Bioelectronics for Full Spherical Electrophysiological Interrogation of 3D Tissues

**DOI:** 10.64898/2026.05.26.728005

**Authors:** Huan Li, Xiaoyu Wang, Yuhang Song, Xin Hu, Jun Yao

**Author notes:** Corresponding author. (J.Y.). These authors contributed equally to this work (Huan Li, Xiaoyu Wang).

## Abstract

Three-dimensional (3D) in vitro tissue models are emerging as powerful platforms for studying development, disease, and therapeutic responses, where close monitoring of electrophysiological activity is essential. However, existing probing methods remain limited in accessibility or spatiotemporal resolution for comprehensive electrophysiological mapping of suspended 3D tissues that closely mimic the native environments. Here we introduce a Venus flytrap-inspired bioelectronic mesh system that enables the full spherical enclosure of 3D tissues in a suspended configuration. The system consists of two hemispherical meshes that envelop the tissue, constructed from highly flexible, stretchable, cell-scale ribbons interconnected into a tissue-compliant network with integrated recording electrode arrays. This architecture enables intimate, conformal tissue integration and supports stable electrophysiological recordings over 300 days. The high-resolution recordings allow precise tracking of local dynamics and correlated global signaling, enabling comprehensive assessment of tissue development as well as detailed evaluation of drug responses for disease modeling. Beyond single tissues, the mesh architecture is extended to fully enclose assembloids composed of multiple tissues, enabling characterization of cross-tissue signaling relevant to advanced heterogenous tissue modeling. Furthermore, the system is translated into array-based platforms, demonstrated by a 4×4 array integrating 1024 electrodes, for high-throughput tissue sampling and cross-study analysis. The developed bioelectronic system and integration method provide a broadly applicable platform to advance electrophysiological studies across diverse tissues and organoids.

## MAIN

In vitro human tissue models are attracting increasing interest to serve as promising platforms for modeling organ development^1,2^, investigating drug mechanisms^3,4^, assessing transplantation strategies^5,6^, and exploring hybrid living systems^7,8^. This trend is further reinforced by recent regulatory initiatives, including the U.S. Food and Drug Administration’s April 2025 plan to phase out animal testing and promote the use of human in vitro models. Compared to traditional planar tissue cultures, three-dimensional (3D) microtissues or organoids can more closely recapitulate cell states, microenvironments, and cell-cell interactions found in the living organs^2,4^, thus gaining further popularity for improved modeling studies. Because electrical signaling plays a critical role in many tissue models, close monitoring of electrophysiological activity is essential for assessing tissue state. Nevertheless, conventional probing techniques for planar tissues face considerable challenges in 3D tissue systems. For instance, optical imaging encounters challenges in accessing 3D morphology due to light scattering and absorption, and its spatiotemporal resolution is often insufficient to resolve fast dynamics across the entire tissue. Electrical recording using planar multielectrode array (MEA), as a gold standard for high-resolution electrophysiology^9^, still faces challenges to effectively access cells in 3D environments.

To this end, several recent advancements have been made to improve MEA recordings in 3D tissue systems. For instance, mesh electronics^10^, composed of an interconnected, ultra-flexible ribbon network, can be co-cultured with cells and progressively integrated into the growing tissue to enable electrophysiological acquisition in a 3D tissue volume^11–15^. While effective, these electronics still rely on a substrate-carrying design with the tissues supported by a planar surface, which differs from the native suspended environment. Additionally, the largely random embedment of mesh network makes it difficult for the spatial registration of electrophysiological activities (e.g., for studying spatiotemporal correlation).

Post-culture integration strategies are also developed as alternative approaches to mitigate the above limitations^16–21^. The general concept is to engineer a hemispherical, soft ‘basket’ to host the 3D tissue for conformal interfacing^17–19^. The strategy offers flexibility to probe tissues at different developmental stages and supports the tissue in a suspended form better mimicking the native environment. Nevertheless, the upholding ‘basket’ design necessitates self-supporting architecture relying on relatively rigid and large structures, yielding structural and mechanical mismatch with tissues. The number of recording sites is also limited (e.g., 16)^20^, constraining spatiotemporal resolution. A recently improved strategy employs a drop-down ‘kirigami’ design to circumvent structural self-support, enabling tissue-like softness for improved tissue interfacing and long-term electrophysiological recordings^21^. However, a common characteristic of these ‘basket’ designs is that they primarily cover only one hemisphere, thereby restricting tissue-wide probing. As a result, high-density, tissue-wide electrophysiological recording in suspended 3D tissue models that mimic the native environment has yet to be achieved.

Here, we demonstrate a bioelectronic mesh system that enables the full spherical enclosure of 3D tissues in a suspended configuration. The constituent ribbons in the mesh achieve cellular feature size and integrate electrode arrays to attain high-resolution recordings across the 3D tissue when implemented in cardiac microtissue. The tissue-like softness in the mesh improves tissue integration and enables record-long electrophysiological probing over 300 days. The full-spherical, high-resolution, and long-term electrophysiological recordings allow not only precise tracking of local dynamics and correlated global signaling for a comprehensive assessment of development but also detailed monitoring of drug effects for disease modeling. The system can be further tailored with a dual-mesh design to fully envelop assembloids for correlation and connectivity study. A 4×4 mesh array integrating 1024 electrodes is demonstrated to show the potential of translating the system into array-based technology for high-efficient assay.

## RESULTS AND DISCUSSION

### Device design and tissue interfacing

We draw inspiration from Venus flytrap (Dionaea muscipula), a carnivorous plant that can close two leaves to capture prey (**Figure 1a(i)**). Our device adopts a similar concept by closing two hemispherical meshes to achieve full spherical tissue enclosure and is therefore termed a *mesh trap* (**Figure 1a(ii)**). To facilitate the transition from a planar lithographic pattern to a hemisphere, the mesh used S-shaped ribbons for high stretchability^22^, following the principle of equal-area partitioning and geometric projection for uniform distribution (**Figure 1b(i)**, Supplementary Fig. 1). The mesh substrate was made from polyimide (PI), chosen for its flexibility, biocompatibility, and convenience with nanofabrication processes^23^. Finite element analysis (FEA) was performed to evaluate the mesh’s mechanical behaviors when enveloping spherical tissues with a typical modulus of 10 kPa^24^ (Supplementary Fig. 2). For the full spherical expansion, FEA revealed an average strain of 0.9% in the mesh ribbons (**Figure 1b(ii)**), which is far below the fracture limit of PI (∼80%)^23^. The average stress over the tissue exerted by the ribbons was 170 Pa (**Figure 1b(iii)**), much lower than values known to induce perturbing mechanical effects, including cell-generated pressures (1–15 kPa)^25^ and the threshold associated with cellular or nuclear deformation (∼5 kPa)^26^.

**Figure 1.**
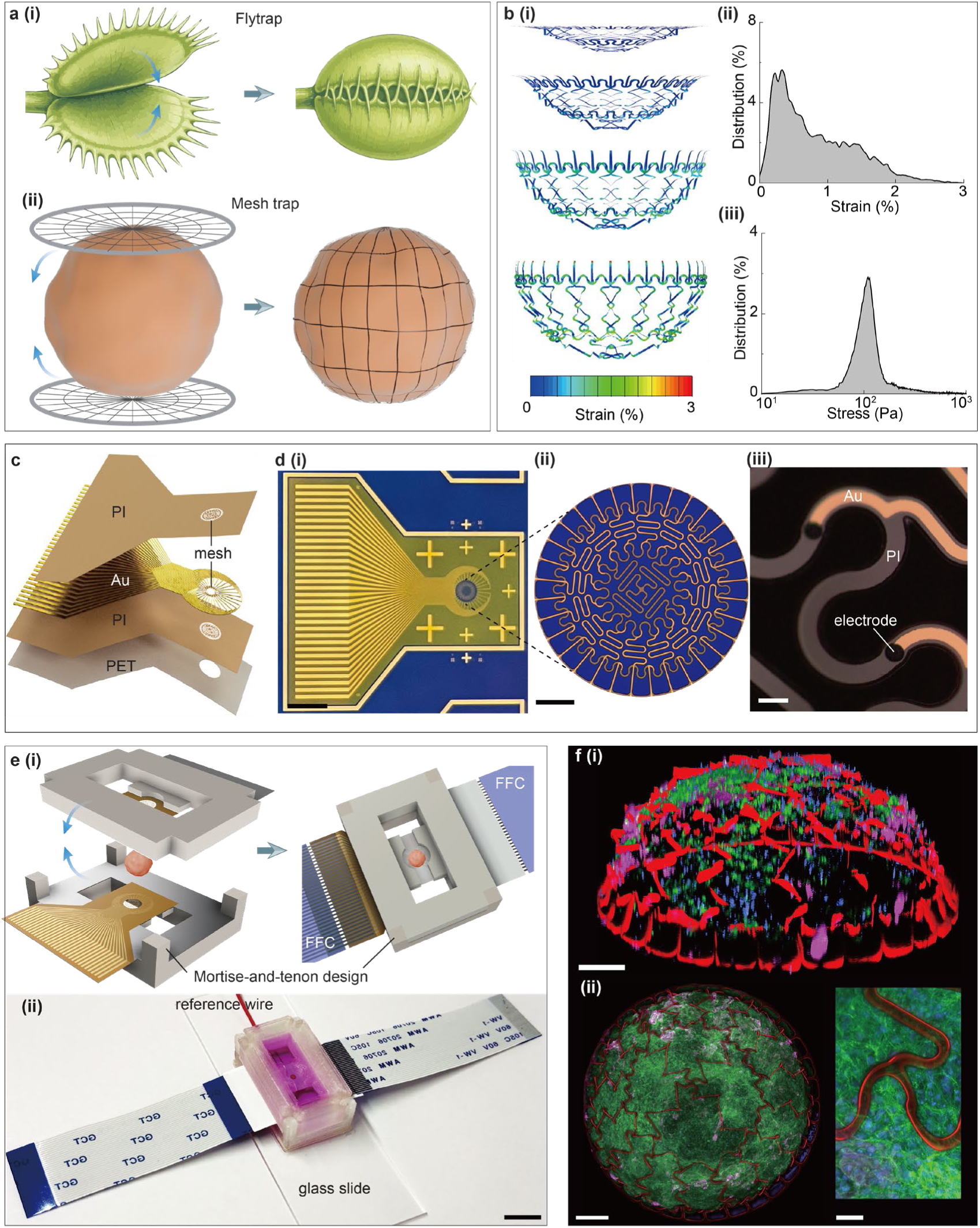
Device design and tissue interface. **a**, Schematics showing the inspiration from (**i**) flytrap for (**ii**) mesh-trap device design. **b**, Mechanical simulations of the (**i**) mesh expansion, (**ii**) its strain distribution and (**iii**) exerted local stress at full expansion. **c**, Exploded-view schematic of the mesh device. **d**, Optical images of (**i**) a fabricated device, (**ii**) its mesh network, and (**iii**) integrated recording electrodes. Scale bars: 3 mm, 0.3 mm, 25 μm. **e**, (**i**) Schematic and (**ii**) photograph of the assembled platform for tissue integration. Scale bar: 8 mm **f**, Confocal fluorescence images of (**i**) a reconstructed 3D view and (**ii**) top view of a tissue (green)-mesh (red) integration, with the inset showing clear surface topology. Scale bars: 0.2 mm, 0.2 mm, 30 µm.

We then translated the design into actual device by photolithographic fabrication (Supplementary Fig. 3). The device featured multilayer structures (**Figure 1c**), with the mesh region covering a circular area of diameter around 1.6 mm, adjustable and comparable to typical microtissue sizes (**Figure 1d(i–ii)**). The constituent ribbons were composed of a thin Cr/Au metal layer (10/100 nm) sandwiched between a top (0.8 µm) and bottom (0.8 µm) PI layers for support and insulation. The ribbon’s resultant thickness of 1.6 µm (Supplementary Fig. 4) and width of 16 µm yielded an ultrahigh flexibility (bending stiffness ∼ 1.4×10^-14^ N·m^2^) order of magnitudes lower than devices with self-support ‘basket’ design^17–19^. 32 recording electrodes with a diameter of 11 µm, further plated with a conductive PEDOT:PSS layer, were integrated on each hemispherical mesh (**Figure 1d(iii)**, Supplementary Fig. 5). Reducing the interconnect feature size could increase the number of recording electrodes to 128 in a hemisphere, resulting in total of 256 for the full sphere (Supplementary Fig. 6). The mesh was surrounded by a non-porous PI substrate for interconnect routing. The entire mesh system was fabricated on a sacrificial layer (e.g., SiO_2_) and subsequently released from the substrate. A polyethylene terephthalate (PET) layer (thickness 0.12 mm), perforated with an opening matching the mesh size, was used to reinforce the mechanical rigidity of the surrounding region for supporting mesh suspension (**Figure 1c**). A culture chamber was 3D-printed using polyethylene terephthalate glycol (PETG) (Supplementary Fig. 7), which is biocompatible^27^ and broadly used in commercial cell culture shake flasks (**Figure 1e(i)**). The chamber consisted of a base for suspending one hemispherical mesh and an upper frame for enclosing the other mesh. A mortise-and-tenon design was used to align the assembly and reinforce a tight seal for hosting culture media. The input/output (I/O) contacts of the meshes remained outside of the chamber for connecting to the recording system via flexible flat cables (FFCs). A platinum wire was integrated into the chamber to serve as reference and ground electrode (**Figure 1e(ii)**). The chamber bottom was made from a transparent glass slide, allowing for convenient optical imaging.

We cultured cardiac microtissues following previous procedures^14^ (Supplementary Fig. 8). We then enclosed a cardiac microtissue in the mesh trap to examine the interfacing morphology. Fluorescence imaging showed that the mesh (red) was spherically expanded to encapsulate the microtissue (**Figure 1f(i)**). The symmetric expansion suggests that the mesh design and assembly strategy can ensure a self-limiting, conformal tissue integration. The mesh ribbons were clearly visible on the surface of the microtissue (**Figure 1f(ii)**), suggesting that they exerted little mechanical perturbation (e.g., local cutting) to the tissue, consistent with FEA analysis.

### Electrophysiological recordings

We continued to investigate the functional integration by performing electrical recordings (Supplementary Fig. 9). Notably, all the 64 integrated electrodes (100% yield) captured periodic extracellular action potentials (eAPs) from cells (**Figure 2a**). Statistics over five integrations yielded consistent high-yield recording (>97%, Supplementary Fig. 10). The high recording yield further supported that the mesh trap formed intimate interface with the microtissue. The captured eAPs featured not only a clear spike signature of the fast upstroke dynamics during depolarization but also a concomitant RT interval related to repolarization (**Figure 2b**). The high temporal resolution (50 μs, 20 kHz sampling) also captured relative delays between the signals (**Figure 2a**, right), revealing the spatiotemporal correlation across the global scale. Altogether, the mesh trap uniquely captured high-resolution local and global electrophysiological dynamics for comprehensive investigation of the relationship to tissue state and development.

**Figure 2.**
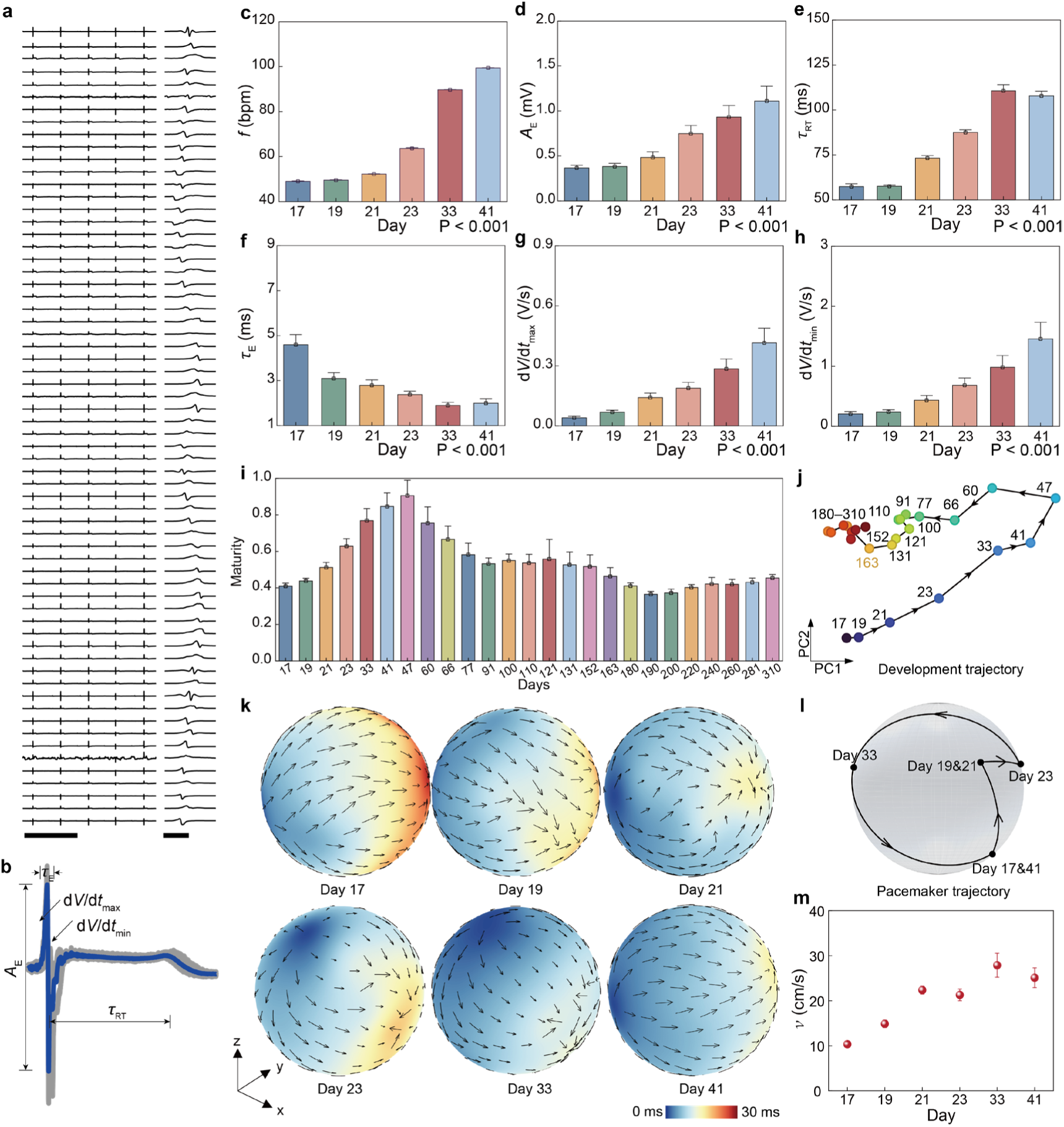
Electrical recordings and developmental analysis. **a**, Representative recordings from the 64 electrodes of a mesh trap and (right) temporally resolved extracellular action potentials (eAPs). Scale bars: 2 s, 10 ms. **b**, Representative superimposed eAP signals with corresponding electrophysiological metrics. The blue curve represents the mean waveform. **c–h**, Temporal evolutions of the six electrophysiological metrics. One-way ANOVA is used with the beginning group as control. **i**, Principal component analysis (PCA) of the electrophysiological features over 310 days (300 days post-integration). The maturity is defined by PC1 that captures the dominant variance. **j**, Developmental trajectory of the electrophysiological phenotype analyzed in the PCA feature space. **k**, 3D mapping of the electrical conduction across the tissue sphere. The color and arrow indicates the temporal delay and local velocity, respectively. **l**, Spatial trajectory of the pacing site over time. **m**, Evolution of the conduction velocity (*v*). Data in (c–i) and (m) are presented as mean ± s.e.m.

We extracted key features from the continuous recordings for analysis. The recordings revealed a converged beating rate across different sites (**Figure 2c**), suggesting efficient intercellular electrical coupling for synchronized contraction^28^. The beating frequency (*f*) showed a steady increase from 49 to 99 beats per minute (bpm). Correspondingly, the eAP amplitude (*A*_E_) also exhibited a consistent increase from 0.4 to 1.1 mV (**Figure ^2^d**), consistent with the progressive growth of Na^+^ channel numbers during maturation^29–31^. The RT interval (*τ*_RT_), known to be related to the continuous K^+^ currents during repolarization^32^, increased from 57 to 110 ms (**Figure 2e**), indicating a prolongation of depolarization through K^+^-channels for counterbalancing the ion influx from increasing Na⁺- and Ca²⁺-channel activities^33^. A close look at the fast dynamics revealed a decrease of the eAP duration (*τ*_E_) from 4.6 to 2.0 ms (**Figure 2f**), consistent with an increasing upstroke velocity or shortening depolarization in maturing cells. The derivatives of the signals (d*V*/d*t*) that are related to ion channel activities^13,32^ are further analyzed. The maximum value (d*V*/d*t*_max_) increased from 0.04 to 0.42 V/s (**Figure 2g**), consistent with an increasing activation rate in the Na^+^ channels in maturing cells. The minimum value (d*V*/d*t*_min_) increased from 0.21 to 1.46 V/s (**Figure 2h**), consistent with an increasing activation rate in the fast K^+^ channels during the initial depolarization.

To better correlate electrophysiological activities to tissue developmental state, we applied principal component analysis (PCA), which is a widely used dimensionality-reduction method that can project the electrophysiological features onto a pair of orthogonal components to capture the dominant sources of variance and reconstruct a simplified developmental trajectory^34^. The first principal component (PC1) encompassing the most total variance, was used as the main composite index to reflect the overall maturity. PC1 increased continuously till day 47 and subsequently decreased (**Figure 2i**). Consistent with this trend, the trajectory analysis in the PCA state space revealed a pronounced inflection point at day 47, after which the trajectory deviated from the initial maturation direction (**Figure 2j**). These results indicate a transition from a steady electrophysiological maturation to an aging-like decline, consistent with trends observed in other cardiac tissues^35^. Notably, the mesh system continuously tracked tissue development for over 300 days (Supplementary Fig. 11), suggesting intimate tissue integration for long-term electrophysiological monitoring. To our knowledge, this is the longest reported recording duration for bioelectronics conformally coupled to 3D human tissue models.

The full-spherical, high-resolution recordings enabled the unique opportunity to study global signaling across the tissue. Combining spatial location and temporal differentiation in the recording signals, a series of 3D conduction maps were reconstructed across different developmental stages. The maps revealed that the electrical conduction consistently originated from a single pacing site and propagated across the microtissue (**Figure 2k**). Interestingly, the pacing site shifted over time (**Figure 2l**), consistent with a continuous remodeling in ion channels and gap junctions that shift the funny current (*I*_f_) responsible for the initial spontaneous depolarization^36^. The high-density recordings further enabled a spatial mapping of local conduction velocity (*v*), revealing a convergent distribution indicative of coordinated development.

The enabled high-resolution mapping can pinpoint aberrant signaling (e.g., wavefront collision and focal activity), thus significantly substantiating disease and dysfunction modeling. The conduction velocity (*v*) exhibited an overall increasing trend by rising from 7 to 30 cm/s (**Figure 2m**), consistent with maturing electrophysiological phenotype. Notably, these values exceed typical values reported in engineered cardiac tissues^37–39^ and fall within the range reported in adult cardiomyocytes^40^. The enhanced conduction, attributed to improved developmental reorganization and gap junction coupling^41^, suggest that the suspended microenvironment provided by the mesh trap may improve electrophysiological development. This attribute, together with advances in full tissue-scale, stable, and high-resolution probing, positions the mesh system as a unique platform for modeling development, studying dysfunction, and investigating connectomics in electroactive tissues.

### Drug models

The growing recognition of the limited translational fidelity of animal models, together with recent regulatory momentum, has made in vitro tissue models important platforms for preclinical drug evaluation^42^. Many drugs target modulating ion-channel activities for their central role in governing physiological functions. By capturing high-resolution local and global electrophysiological dynamics, the mesh system can enable more comprehensive assessment of drug effects. We therefore used the mesh trap to probe different drug effects from lidocaine, E-4031, and isradipine (**Figure 3a**, Supplementary Fig. 12), which are blockers to the three prominent Na^+^, K^+^, and Ca^2+^ channels, respectively^32,43^.

**Figure 3.**
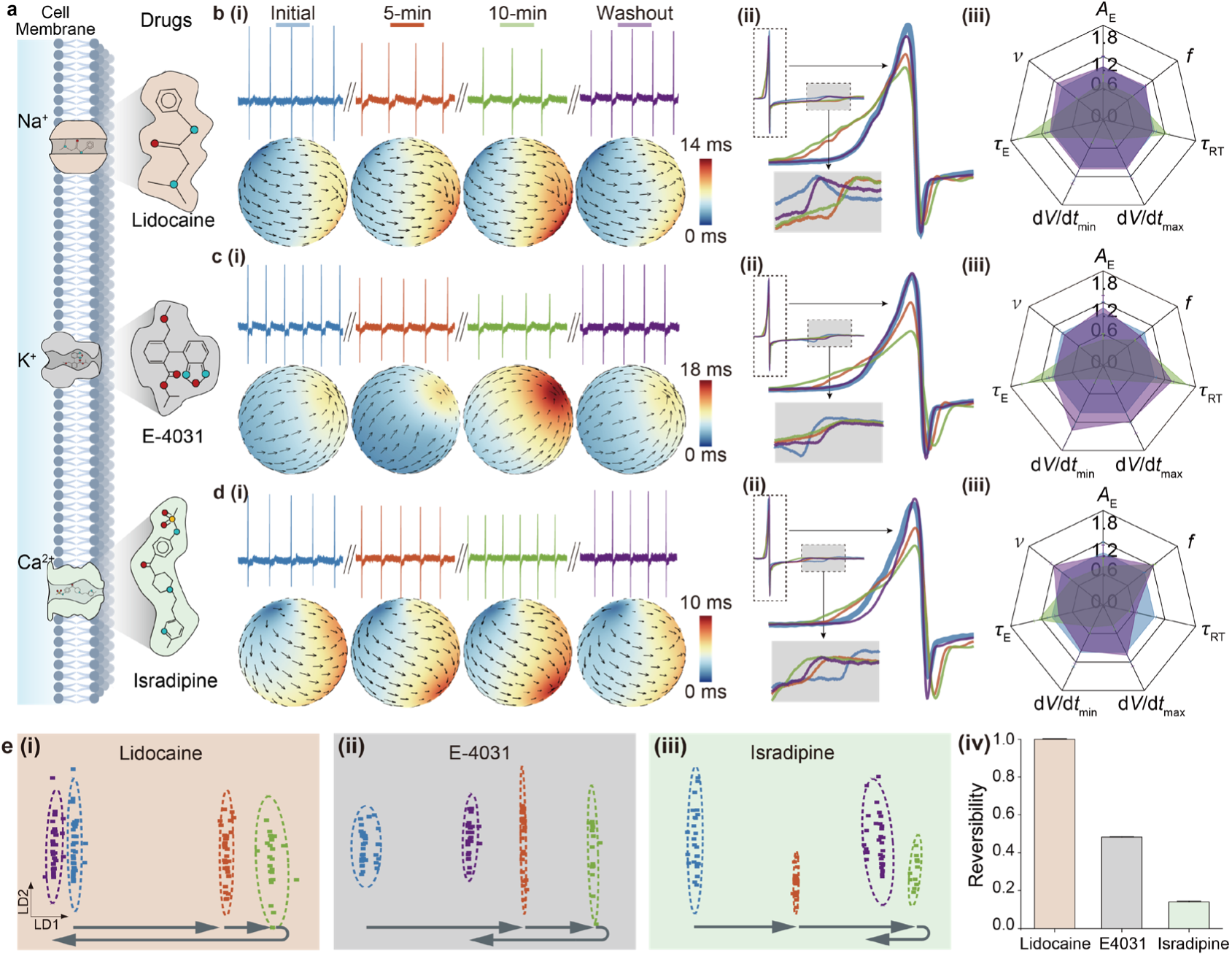
Drug effect analysis. **a**, Schematics of three drugs targeting different ion channels. **b–d**, Evolutions of electrophysiological activities in tissues administrated with Lidocaine, E-4031, and Isradipine, respectively. These include: (**i**) Representative recordings and conduction maps at baseline, 5 and 10 minutes after drug administration, and following washout, respectively; (**ii**) Representative superimposed waveforms at corresponding stages; (**iii**) Radar plots summarizing the evolutions of normalized electrophysiological metrics. **e**, (**i**-**iii**) Evolutions of electrophysiological metrics in the feature space by linear discriminant analysis (LDA). Each color denotes a stage same as that in (b–d). (**iv**) Corresponding drug reversibility based on the LDA trajectories. Data in b–d(iii) and e(iv) are presented as mean ± s.e.m.

Representative recordings showed that both the eAP amplitude (*A*_E_) and frequency (*f*) decreased after lidocaine (20 µM) administration (**Figure 3b(i)**, upper panel). The global signaling, uniquely captured by the mesh, exhibited a trend of increasing delay (bottom panel), suggesting a reduction in conduction velocity (*v*). These features largely recovered after drug washout. Detailed analysis revealed a 30% reduction in *A*_E_ (**Figure ^3^b(ii–iii)**), consistent with the role of lidocaine in blocking Na^+^ influx that is mainly responsible for eAP signal. The slowing in *f* and *v* is further connected with depolarization details captured in fast dynamics. Specifically, eAP duration (*τ*_E_) was prolonged by 53%, accompanied by the reduction of d*V*/d*t*_max_ (58%) and d*V*/d*t*_min_ (63 %) (**Figure 3b (iii)**), suggesting the reduction of upstroke and downstroke velocity during depolarization. As the upstroke velocity is positively related to conduction velocity^30,44^, the reduction in *v* (from 34 to 19 cm/s) was consistently observed. Meanwhile, the slowing Na^+^ dynamics can lead to delay in subsequently coupled Ca²⁺ and K⁺ currents^45,46^ to cause repolarization delay, reflected by a RT interval (*τ*_RT_) prolongation (25%) and a concomitant reduction in *f* (29%) from the recordings.

Drug E-4031 (50 nM) induced similar trends in *A*_E_, *f*, and conduction delay with those by lidocaine (**Figure 3c(i)**). This is because the blockage of K⁺ channels elevates resting membrane potential and thereby downregulates Na⁺-channel activity^44^, causing similar effect to the direct blockage of Na^+^ channels by lidocaine. Nevertheless, as E-4031 directly reduces K^+^-channel activity and thus significantly delays repolarization, its effect on *τ*_RT_ prolongation (62%) was more prominent (**Fig. 3c(ii–iii)**). Isradipine treatment (20 nM) produced similar electrophysiological trends in *A*_E_, *f*, and conduction delay (**Figure 3d(i)**), because the Ca²⁺-channel blocker is known to have a secondary effect in suppressing Na^+^-channel current^44^. However, isradipine generated opposite trends of *τ*_RT_ shortening (51%) and *f* increase (25%) (**Figure 3d(ii–iii)**), consistent with the mechanism that a reduced inward Ca²⁺ current in turn shortens the depolarization involving the counterbalancing outward K^+^ current.

The above results and analysis demonstrated that the mesh trap captured multi-dimensional spatiotemporal electrophysiological features encompassing both shared characteristics and distinct variations. To better distinguish drug-specific effects and development, we further applied linear discriminant analysis (LDA). In the LDA space, the three drugs exhibited distinct trajectories spanning the initial, 5-min, 10-min, and washout phases (**Figure 3e(i–iii)**), with the degree of looping overlap in the trajectory indicating the level of reversibility (**Fig. 3e(iv)**). Lidocaine exhibited near-complete recovery (**Figure 3e(i)**), consistent with its highly reversible electrostatic binding to Na⁺ channels, which is modulated by the competing channel conformational changes during depolarization and repolarization^47^. E-4031 showed partial recovery (**Figure 3e(ii)**), which was attributed to the trapping of drug molecules within the closed channels upon inactivation, thereby hindering unbinding during washout^48^. Isradipine displayed relatively low recovery (**Figure 3e(iii)**), consistent with its slow washout kinetics resulting from its high lipophilicity^49^. For all three drugs, the signaling across the tissue remained as a coherent wavefront, suggesting preserved cell-cell electrical coupling despite modulation of ion-channel activities (**Figure 3b–d(i)**, bottom panels).

### Dual-mesh traps for assembloids

While the above investigations have demonstrated the mesh’s unique ability in capturing comprehensive electrophysiological activity in individual tissues for developmental and drug studies, recent trend shows a strong interest in merging tissues for assembloids to enhance structural and functional integration for improved modeling^50–52^. Although some previous platforms demonstrated the probing of electrophysiological activities in assembloids^17,21^, they fell short of a full-coverage and high-density interrogation. Here, we constructed a dual-mesh trap system that can extensively probe electrophysiology in assembloids for studying correlated dynamics.

The dual-mesh system merged two neighboring meshes (**Figure 4a**), with each following the design of the previous individual mesh (**Figure 1d**). The two meshes shared an arc ∼ 60°, a typical interfacial portion in assembloids^53^. Each hemispheric mesh integrated 28 recording electrodes, resulting in a total number of 104 for the full enclosure. The resultant dual-mesh design is fully compatible with the single-tissue chamber architecture and can employ the same mortise-and-tenon encapsulation strategy (**Figure 4b**).

**Figure 4.**
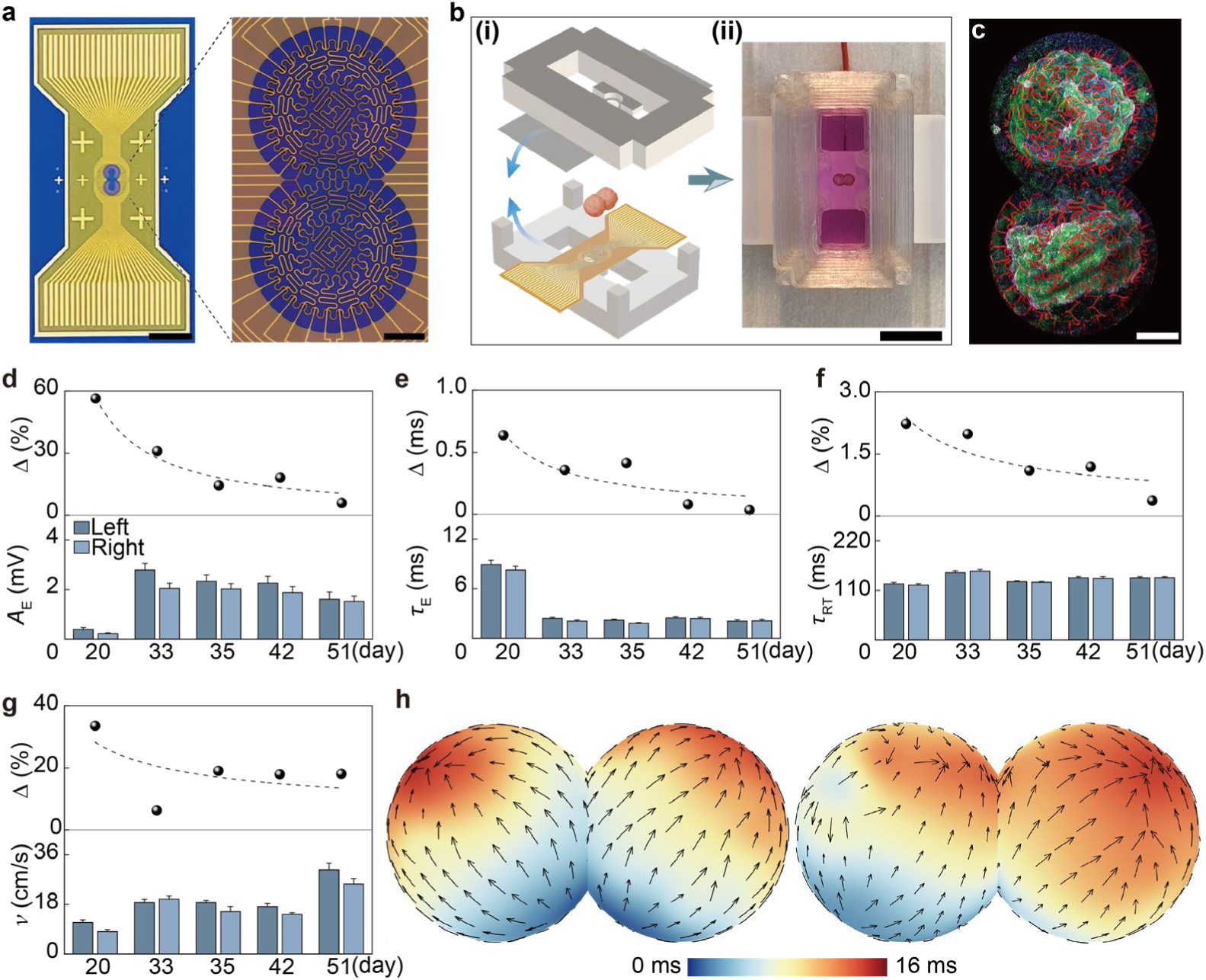
Dual-mesh trap for assembloids. **a**, Optical images of a fabricated dual-mesh. The right is a stitched image. Scale bars: 4 mm, 40 μm. **b**, (**i**) Schematic and (**ii**) photograph of an assembled dual-mesh platform for tissue integration. Scale bar: 8 mm. **c**, Fluorescence image of an assembloid (green)-mesh (red) integration. Scale bar: 0.3 mm. **d–g**, (Bottom) evolutions of eAP amplitude (*A*_E_), duration (*τ*_E_), RT interval (*τ*_RT_), and conduction velocity (*v*) in the left and right microtissues and (top) their difference. **h**, Representative 3D conduction maps featuring (left) bidirectional and (right) unidirectional propagation. Data in (d–g) are presented as mean ± s.e.m.

We employed cardiac assembloids to demonstrate integration and functional characterization, which carried practical relevance because cross-tissue signaling underpins successful cardiac grafting for potential transplantation. Two cardiac microtissues initially cultured in separate environments were integrated into the dual-mesh trap. Fluorescence imaging showed that the two microtissues abutted each other to form an assembloid, which also established a conformal interface with the mesh trap (**Figure 4c**). Notably, 102 of the 104 recording electrodes (yield > 98%) successfully detected electrical eAPs (Supplementary Fig. 13), further confirming the intimacy in mesh-tissue morphological and functional integration.

We used recorded high-resolution electrophysiological activities to investigate the functional interaction and evolution in the assembloid. The assembloid demonstrated a steady increase in beating frequency *f* (Supplementary Fig. 13c), consistent with the trend observed in individual microtissues (**Figure 2c**). The two microtissues maintained converged contraction, suggesting a close electrical coupling. The eAP amplitude (*A*_E_) showed a substantial increase on day 33 and then maintained a stable level (**Figure 4d**, bottom panel), while its duration (*τ*_E_) showed an opposite trend (**Figure 4e**, bottom panel). These trends are consistent with maturing phenotypes and timespan, reflecting the increasing Na^+^-channel activity for higher and faster Na^+^ influx. *A*_E_ and *τ*_E_ between the two microtissues showed consistent and continuous convergence, with the difference decreased from 56% to 6% and 0.64 ms to 0.03 ms, respectively (**Figure 4d, e**, top panels). The RT interval (*τ*_RT_), primarily related to K^+^ channel activity, while maintaining an overall stable level, exhibited a converging trend between the two microtissues with their relative difference decreased from 2.2% to 0.4% (**Figure 4f)**. These coordinated convergences are consistent with a ‘tuning’ effect from synchronized signaling, in which ion channels were reported to have ‘molecular memory’ with their availability shaped by prior activation history and pacing frequency^54,55^.

We further analyzed the spatiotemporal signaling dynamics in the assembloid. The conduction velocity (*v*) showed an overall increase and a general trend of convergence but retained some spatial discrepancy (15%) between the two microtissues (**Figure 4g**, bottom panel). As *v* depends on both Na⁺-channel activity and cytoplasmic and gap junctional resistances^30,44^, the observed convergence in Na⁺ channel activity suggested by eAP features (**Figure 4d, e**) indicates that tissue resistance predominantly accounts for *v* discrepancy. From the reconstructed 3D conduction map uniquely enabled by the full-enclosure integration, we observed two distinct propagation modes over the course. One exhibited a bidirectional mode with the signaling bifurcating from a pacing site at the joining border (**Figure 4h**, left). The other featured a unidirectional one with the signaling advancing from the distal side of one tissue to the other (**Figure 4h**, right). In the current assembloid without functional differentiation, the interchangeable wave-like pacing modes, similar to those observed in the individual tissues, indicate close functional integrity between the components. The design can be extended to accommodating more microtissues in an assembloid (Supplementary Fig.14). The platform can be readily applied to multi-chamber cardioids with functional differentiation for high-resolution quantification of intercompartmental signaling^52^. It can also be extended to other heterogeneous assembloids, including those incorporating blood vessel organoids to study vascularization effect and neural assembloids to investigate circuit development.

### Array platform

The efficiency in many bioassays is underpinned by array-based technology for parallelism and efficiency. We translated the mesh system into an array platform, demonstrated with 2×2 (Supplementary Fig. 15), 2×4 (Supplementary Fig. 16), and 4×4 arrays (**Fig. 5a**) compatible with academic wafer-scale fabrication (**Figure 5b(i)**). A total number of 1024 electrodes were integrated in the 4×4 array. On each side, two 64-channel FFCs were used for signal routing (**Figure 5b(ii)**). Culture chambers with corresponding arrays of openings were fabricated by 3D printing, using similar mortise-and-tenon design between the upper and lower frames for mesh alignment and fixation (**Figure 5b(iii)**).

**Figure 5.**
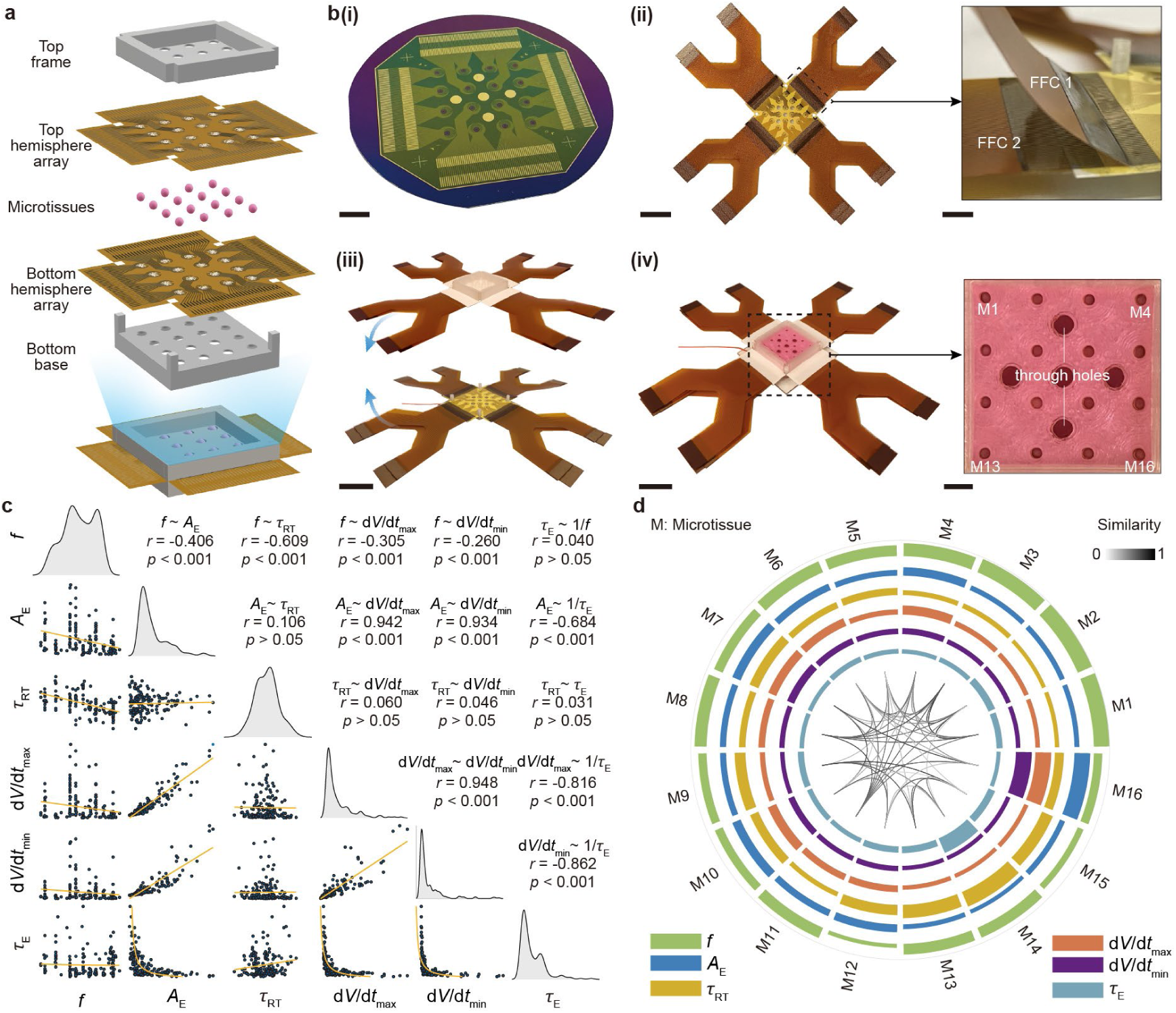
Mesh-trap array. **a**, Exploded view of the assembled array system. **b**, Photo images of (**i**) a 4×4 mesh array fabricated on wafer, (**ii**) the released array connected with flexible flat cables (FFCs), each side having bilayer connections (right), (**iii**) a pair of upper and lower mesh arrays for system assembly, and (**iv**) an assembled array system, (right) integrating cardiac microtissues (M). Scale bars: 8 mm, 25 mm, 10 mm, 35 mm, 25 mm, 5 mm. **c**, Pairwise correlation matrix of electrophysiological metrics across different microtissues. The diagonal panels show the distributions of individual metrics and the off-diagonal panels display pairwise relationships with the Spearman correlation coefficients (*r*) and statistical significance (*p*). **d**, Circos map of electrophysiological phenotypes and pairwise similarity between the 16 microtissues in the 4×4 array. Each sector corresponds to a microtissue (M). The concentric rings represent different electrophysiological metrics. The connecting arcs suggest pairwise similarity, with darker color indicating higher similarity. *p* is calculated using two-sided Spearman rank correlation.

We integrated 16 cardiac microtissues into a 4×4 array to evaluate parallel electrophysiological characterizations (**Figure 5b(iv)**), with five through-holes in the culture chamber added to improve fluid exchange between the upper and lower compartments. Pairwise correlation analysis was employed to reveal the distribution and correlation in the electrophysiological phenotypes across all microtissues (**Figure 5c**). The diagonal distributions illustrated the overall variability of each parameter across microtissues. The Spearman correlation coefficient (*r*) was used to quantify the strength of pairwise relationships, with a larger absolute value indicating stronger correlation. *A*_E_, d*V*/d*t*_max_, and d*V*/d*t*_min_ were tightly coupled (*r* = 0.93–0.95), and inversely scaled with the eAP duration (*τ*_E_) (|*r*| = 0.68–0.86), reflecting the convergence in Na^+^- and K^+^-channel activities that govern rapid depolarization dynamics. Meanwhile, RT interval (*τ*_RT_) was statistically decoupled from depolarization-related metrics (|*r*| < 0.11), indicating a functional dissociation between the repolarization and depolarization processes. The beating frequency (*f*) exhibited moderate correlations with other parameters (|*r*| = 0.26–0.41) but a stronger association with *τ*_RT_ (|*r*| = 0.61), consistent with the role of repolarizing currents in setting pacemaker rate. By contrast, *f* exhibited little correlation with the *τ*_E_ (*r* = 0.04), indicating that pacing frequency is largely decoupled from depolarization timescale.

We constructed a multidimensional Circos phenotypic map incorporating these electrophysiological metrics for inter-tissue comparison (**Figure 5d**). The sectors denote individual microtissues, concentric rings encode the relative magnitudes of electrophysiological metrics, and the greyscale of connecting cords represent pairwise similarity of collective electrophysiological phenotypes. The connecting cords showed closer distribution compared to individual metrics, indicating that the integrated tissue array maintained reasonable uniformity in the collective phenotypes.

The above recordings and analysis show that the mesh array can provide efficient tissue sampling and cross comparison for various studies related to genetic, developmental, and pharmacological effects. Though not demonstrated here, array platforms for the dual-mesh system (Supplementary Fig. 17) can be similarly fabricated to expand the applicability to assembloid arrays.

## CONCLUSION

This work presents a bioinspired mesh system that enables full spherical enclosure of 3D tissues in a suspended configuration. The cell-level feature size and tissue-level softness in the mesh establish an intimate, conformal tissue interface that supports long-term stable electrophysiological interrogation. The integrated high-density recording electrodes enables high-resolution spatiotemporal analysis of electrophysiological phenotypes and reconstruction of 3D signaling maps, providing a more comprehensive assessment of tissue development and drug response. Beyond single tissues, the mesh architecture can be extended to fully envelop assembloids composed of multiple microtissues, enabling characterization of cross-tissue signaling relevant to advanced models involving heterogenous tissue integration. The mesh system can be translated into array-based platforms for high-throughput tissue sampling, cross-validation, and control studies, which can further synergize with data-driven and AI-assisted analytics for improved precision and efficiency.

## METHODS

### Mesh design

The mesh design followed principles of equal-area partitioning and geometric projection as described in Supplementary Fig. 1. Briefly, the spherical surface was first subdivided into 64 equal-area regions in latitude-longitude coordinates. This approach can be extended to generate a larger number of channels if needed. The geometric center of each region was then calculated. Adjacent center points were connected, with the corresponding arc lengths (*L*₁) computed. Then these arcs were projected onto the XY plane, with the corresponding planar distances (*L*₂) calculated. *L*₂ was replaced by an S-shaped curve, stretchable to original arc length *L*₁ during expansion. The ratio between the curved *L*_2_ and *L*_1_ defines a correction factor. Mesh designs with correction factors of 1, 1.25, 1.5, and 1.75 were evaluated by mechanical simulations, revealing the optimal result by using a factor of 1.75 (Supplementary Fig. 2). Corners at the joints were rounded to improve mechanical robustness^21^.

### Simulation analysis

Finite element simulations were used to evaluate the mechanical interaction between the mesh and microtissue (Abaqus/CAE 2023, Dassault Systemes). Owing to the geometric symmetry and to reduce computational cost, the simulation was performed in a hemisphere. The microtissue was prescribed to move downward against the mesh. The mesh device was discretized by triangular surface meshes with a minimum element size of 6.3 µm. The microtissue was discretized by tetrahedral volumetric meshes with a minimum element size of 7 µm. A typical Young’s modulus of 2.5 GPa, density of 1.42 g/mL, and Poisson’s ratio of 0.34 were employed in the polyimide (PI) material. A density of 1.0 g/mL and Poisson’s ratio of 0.5 were used for the tissue. Initial simulations took the microtissue as a rigid sphere to find the optimal correction factor for mesh expansion with economic computational power (Supplementary Fig. 2). After that, a more detailed simulation was conducted by using a spherical tissue with bio-realistic mechanical properties (Figure 1b). A transient dynamic approach was adopted to avoid convergence challenge encountered in static analysis. Specifically, the mesh was first stretched with the displacement held. Then system was allowed to progressively relax toward a quasi-equilibrium under this constrained state. During the relaxation, minimized kinetic energy was used as indicator of reaching a stable state, which is conceptually analogous to an annealing process. Strain was quantified using the maximum in-plane principal strain to capture the dominant deformation mode of the ultrathin mesh under plane-stress conditions. Stress was assessed using the von Mises equivalent stress, which provides a scalar measure of multiaxial stress in ductile materials.

### Bending stiffness

The bending stiffness of the PI ribbon in the mesh was estimated using classical Euler–Bernoulli beam theory, taking the geometric parameters of a width (*b*) 16 μm and thickness (*h*) 1.6 μm. The bending stiffness was calculated as *EI* = *Ebh*^3^/12, where *E* is the Young’s modulus of PI, yielding *EI* ∼ 1.4 ×10^-14^ N·m^2^. The ultrathin metal layers (10 nm Cr and 100 nm Au) were not considered, as they contributed negligibly to the overall stiffness.

### Device fabrication and system assembly

Device fabrication was performed using standard microfabrication as illustrated in Supplementary Fig. 3, which consists of the following steps. (1–3) A polyimide (PI) precursor (Sigma-Aldrich, 575771) was spin-coated (6000 rpm) on a silicon substrate covered with a layer of 300-nm SiO₂ (Nova Electronic Materials) and baked at 180 °C for 5 min. A layer of photoresist (S1813, Kayaku Advanced Materials) was subsequently spin-coated (2000 rpm) and baked at 115 °C for 5 min. (4–6) Photolithography was performed to define the mesh pattern in the bottom PI layer, with the photoresist layer subsequently removed by acetone. The patterned PI layer was then cured at 250 °C for 30 min. (7–11) Metal interconnects and electrodes were defined by standard lithography (S1813), metal deposition (Cr/Au, 10/100 nm), and lift-off processes. (12) A top passivation PI layer was patterned following similar steps 2–6. (13–16) The fabricated mesh device was picked up by a water-soluble tape, assisted by HF etching of the underlying SiO_2_ layer, transferred onto a polyethylene terephthalate (PET) substrate, and released by dissolving the tape in water. (17) The recording electrodes were electroplated with a PEDOT:PSS layer (Supplementary Fig. 5) using an electrochemical workstation (CHI400C, CH Instruments).

The mesh device was assembled in a customized chamber consisting of a base and top frame, which were 3D-printed using PETG with structural parameters listed in Supplementary Fig. 7. The base was sealed to a glass slide using polydimethylsiloxane (SYLGARD 184). The assembled chamber was sterilized by UV irradiation (30 min). Then the mesh surface was coated with a thin layer of Matrigel and incubated (30 min), with the excess Matrigel gently removed. The pre-cultured microtissue was then transferred into the mesh cavity using a pipette tip. The top frame was mounted onto the base to complete the assembly. A biocompatible UV adhesive was used to provide auxiliary fixation. The mesh-tissue integration was subsequently maintained in a cell culture incubator (Vios 160i, HERACELL) at 37 °C with 5% CO₂.

### Cell differentiation and tissue culture

Human pluripotent embryonic stem cells (hPESCs; WAe009-A/WA09, H9; WiCell; NIH registration no. 0062, female) were expanded under feeder-free conditions on Matrigel-coated 60-mm dishes (10 µg mL⁻¹; Corning, CB-40234) in Essential 8 medium (Gibco, A1517001) prepared in DMEM/F-12 (Gibco, 11-320-033), with passaging every 3–4 days. Vendor-provided authentication and quality control (including morphology assessment, isoenzyme profiling, and mycoplasma testing) were used. Cardiomyocyte differentiation was induced by temporal modulation of Wnt/β-catenin signaling using a small-molecule protocol^14,43,56^. Briefly, hPESCs were seeded in 12-well plates and differentiated upon reaching 90–100% confluence (typically 2–3 days). On day 0, the medium was changed to RPMI 1640 (Gibco, 11-875-101) supplemented with 1% B27 minus insulin (Gibco, A1895601; RPMI/B27-insulin) and CHIR99021 (8 μM; Tocris, 252917-06-9) for 24 h. On day 1, cultures were maintained in RPMI/B27-insulin. On day 3, IWR-1-endo (5 μM; Tocris, 1127442-82-3) was added in RPMI/B27-insulin; the inhibitor was removed on day 5 with a return to RPMI/B27-insulin. On day 7, cultures were switched to RPMI 1640 containing 1% B27 (with insulin) (Gibco, 17-504-044, RPMI/B27+insulin). Media were refreshed every other day thereafter. Contractile activity typically emerged around days 7–9, and cells were harvested for 3D microtissue culture during days 7–12.

Microtissue was formed following a previously reported 96-well microplate-based workflow^57^ (Supplementary Fig. 8). Briefly, cardiomyocytes were rinsed once with 1× DPBS to chelate Ca²⁺ and minimize contraction during dissociation. Each well was incubated with 0.5 mL of 0.05% trypsin-EDTA (Gibco, 25300054) for 4–5 min at 37 °C. After aspirating the enzyme solution, cells were gently triturated in 2 mL RPMI 1640 containing 1% B27 using a 1 mL pipette tip. The suspension was collected into a 15 mL conical tube, centrifuged (250× g, 3 min), and resuspended in RPMI 1640 + 1% B27 supplemented with 10 µM Y-27632 ROCK inhibitor (Tocris, 12-541). Single-cell suspensions were dispensed at 400000–500000 cells per well in round-bottom 96-well plates and briefly centrifuged (250× g, 3 min) to promote aggregation. Within 2–3 days, aggregates compacted into spheroids and detached from the well bottom. The spheroids were transferred to 24-well plates, where beating gradually recovered over the following 3–4 days and the tissues matured into spherical microtissues. Microtissues were maintained in RPMI/B27+insulin with daily medium renewal and were subsequently transferred onto mesh devices for various characterizations. Microtissues used for single-mesh, dual-mesh, and array testing were integrated with devices at 10, 14, and 19 days after differentiation, respectively.

### Whole-mount immunostaining and confocal imaging

Whole-mount immunostaining was performed in microtissues following our previous work^14^. Briefly, tissue samples were rinsed three times with DPBS and fixed in 4% paraformaldehyde in DPBS at 4 °C overnight. After three washes with PBS, samples were incubated in 30% (w/v) sucrose in DPBS at 4 °C for 24–48 h. Samples were then washed with DPBS (3×) and permeabilized in 0.2% Triton X-100 (Thermo Fisher Scientific) at 4 °C overnight. Following DPBS washes (3×), samples were blocked in 10% normal goat serum for 2 h at room temperature and incubated with anti-cardiac troponin T (cTnT; 1:250; Invitrogen, MA5-12960) in blocking buffer at 4 °C overnight in the dark. Samples were washed with DPBS (2×, 15 min each) and incubated for 2 h at room temperature with Alexa Fluor 647 goat anti-mouse secondary antibody (1:500; Invitrogen, A-21235), DAPI (1:1000; Invitrogen, D1306), and Alexa Fluor 488 phalloidin (1:50; Invitrogen, A12379) diluted in blocking buffer. After PBS washes (2×, 2 h each), samples were mounted with Fluoromount-G for microscope imaging (A1R Confocal Microscope, Nikon). Mesh (red), cytoskeleton (green), cardiac-specific protein (purple), nuclei (blue) are shown.

### Electrical measurements

Electrical recordings were performed using an Intan RHD recording system (Supplementary Fig. 9). A customized printed circuit board (PCB) converter was used to connect the flexible flat cable (FFC) to the recording headstage. Signals were acquired at a sampling rate of 20 kHz. Platinum wires (99.99% purity) were used as reference and ground electrodes. All measurements were conducted within a home-built electromagnetic shielding enclosure mounted on a vibration-isolated table.

### Drug assays

Lidocaine (20081), E-4031 (15203), and isradipine (17536) were purchased from Cayman Chemical. All compounds were first dissolved in dimethyl sulfoxide (DMSO), a solvent has negligible effect on microtissue^14^, and subsequently diluted into the culture medium, resulting in a final DMSO concentration below 0.1%. Prior to drug application, the tissue was changed with fresh culture medium and allowed to equilibrate in an incubator for 4 h. Recordings acquired after this equilibration period were defined as the initial state. Following a 15-min baseline period, drugs were added, and measurements were performed after 5 min and again after an additional 5 min. Subsequently, the drug was washed out by drug-free culture medium, and a final recording was performed to assess recovery. Raw results are presented in Supplementary Fig. 12.

### Statistical analysis

Data preprocessing: Raw recording data (rhd format) were converted into MATLAB-readable files using Intan-MATLAB scripts. Custom-written MATLAB code was used for bandpass filtering and removal of 60 Hz power-line noise. Waveforms (e.g., eAP) from the same channel were superimposed and averaged to obtain a mean value, from which electrophysiological metrics were extracted. d*V*/d*t*_max_ and d*V*/d*t*_min_, corresponding to the steepest slopes on each side of the depolarization peak, were obtained by smoothing the waveform using a Savitzky-Golay filter followed by numerical differentiation. The eAP duration (*τ*_E_) is defined as the time interval between the two. Extracted parameters were manually inspected to ensure accuracy. Channels with signal features not reliably identifiable were excluded from subsequent analyses.

Principal component analysis (PCA): Extracted electrophysiological metrics were standardized using z-score normalization. PCA was performed using a NIPALS-based algorithm, involving iterative extraction of dominant components without decomposition of the covariance matrix. The first principal component (PC1), capturing the main variance in the feature space, was interpreted as electrophysiological maturity. The developmental trajectory was constructed by plotting the centroid of each channel in the PCA space defined by PC1 and PC2.

Linear discriminant analysis (LDA): Extracted electrophysiological metrics related to drug tests were standardized by z-score transformation. LDA was applied to these data to extract the discriminant direction. The first two discriminant functions (LD1 and LD2) were used for visualization. The reversibility was quantified by using LD1.

3D conduction mapping: The local activation time (LAT) was determined by the minimum derivative of each eAP signal, which is a widely accepted marker of electrical activation. The collected discrete LATs were transformed into a continuous activation-time field using radial basis function interpolation formulated on spherical geodesic (great-circle) distances.

Conduction velocity: Conduction velocity (*v*) was computed from the reconstructed spatial distribution of LATs. Specifically, neighboring LATs were used to fit a local plane. *v* was then defined as the inverse of the LAT gradient. To ensure reliable estimates, local gradient fitting was weighted by spatial proximity (e.g., close points contribute more than distal ones). Gradient estimation was further performed using regularized regression to reduce sensitivity to noise and spatial irregularity. In assembloid *v* reconstruction, each microtissue was modeled as a unit sphere positioned within a shared 3D coordinate system based on the experimental layout. Discrete LATs from both tissues were mapped onto a complete two-sphere channel configuration. A single global radial basis function was used to reconstruct the LAT field from all valid channels. To avoid unrealistic coupling across the tissue interface, a cross-surface distance penalty reduced inter-sphere influence without fully decoupling them. Conduction velocity was then computed separately for each microtissue using local tangent-plane gradients, ensuring estimates relied only on within-sphere neighborhoods despite global interpolation.

Pairwise coupling matrix analysis: Pairwise relationships between extracted electrophysiological metrics were visualized using a scatter-correlation matrix. Raw data points were plotted in the lower triangle, whereas the diagonal panels showed the corresponding probability density distributions. Pearson’s correlation coefficient (*r*) and two-sided *p* values were computed for each metric pair. To reduce visual clutter, the upper triangle displayed only the relationship label together with *r* and binned *p*. For metric pairs involving eAP duration (*τ*_E_), an inverse regression model (*y* = a·(1/*x*) + b) was applied; otherwise, a linear model (*y* = a·*x* + b) was used. Regression parameters were estimated by least-squares fitting.

Circos-like phenotypic map: Extracted electrophysiological metrics were averaged in each microtissue and further standardized across all tissues. The pairwise phenotypic similarity was quantified using cosine similarity. The metric magnitudes were normalized to ring thickness. Phenotypic similarity was visualized by a connecting chord, with its thickness and transparency scaled by similarity strength.

All data processing and statistical analyses were conducted in MATLAB R2024a, Microsoft Excel, and OriginPro 2025.

## Data availability

The data that support the plots within this paper and other findings of this study are available from the corresponding authors upon reasonable request.

## Acknowledgements

We thank Prof. Alfred Crosby for providing access to computational resources (Abaqus 2023) that supported this project. J.Y. acknowledges support from the Army Research Office W911NF261A034, and National Science Foundation (NSF) 2345733. J.Y. also acknowledges support from National Institutes of Health (NIH) R21EB030216, R33HL175683, NSF 2326703, and Alfred P. Sloan Foundation (FG 2022-18452).

## Author contributions

H.L. and J.Y. conceived the project and designed experiments. H.L. performed experiments in mechanical simulation, device design, fabrication, characterization, chamber design, fabrication, imaging, and in-vitro electrical recordings, MATLAB programming, data analysis, plotting and helped with the cell culture. X.W. performed experiments in cell culture and helped with device fabrication, characterization, imaging, and plotting. Y.S. helped with data analysis. X.H. helped with simulation. H.L. and J.Y. wrote the paper. All authors discussed the results and implications and commented on the manuscript.

## Competing interests

There are no conflicts to declare.

## Notes

### Competing Interest Statement

The authors have declared no competing interest.

## REFERENCES

1 Lancaster, M. A. et al. Cerebral organoids model human brain development and microcephaly. Nature 501, 373–379 (2013).

2 Kim, H., Kamm, R. D., Vunjak-Novakovic, G. & Wu, J. C. Progress in multicellular human cardiac organoids for clinical applications. Cell Stem Cell 29, 503–514 (2022).

3 Harary, I. & Farley, B. In vitro studies of single isolated beating heart cells. Science 131, 1674–1675 (1960).

4 Richards, D. J. et al. Human cardiac organoids for the modelling of myocardial infarction and drug cardiotoxicity. *Nat*. Biomed. Eng. 4, 446–462 (2020).

5 Aoyama, J. et al. Flexible nanoelectronics reveal arrhythmogenesis in transplanted human cardiomyocytes. Science 390, eadw4612 (2025).

6 Revah, O. et al. Maturation and circuit integration of transplanted human cortical organoids. Nature 610, 319–326 (2022).

7 Fu, S. et al. Constructing artificial neurons with functional parameters comprehensively matching biological values. Nat. Commun. 16, 8599 (2025).

8 Michas, C. et al. Engineering a living cardiac pump on a chip using high-precision fabrication. Sci. Adv. 8, eabm3791 (2022).

9 Spira, M. E. & Hai, A. Multi-electrode array technologies for neuroscience and cardiology. Nat. Nanotechnol. 8, 83–94 (2013).

10 Hong, G., Yang, X., Zhou, T. & Lieber, C. M. Mesh electronics: a new paradigm for tissue-like brain probes. Curr. Opin. Neurobiol. 50, 33–41 (2018).

11 Dai, X., Zhou, W., Gao, T., Liu, J. & Lieber, C. M. Three-dimensional mapping and regulation of action potential propagation in nanoelectronics-innervated tissues. Nat. Nanotechnol. 11, 776–782 (2016).

12 Li, Q. et al. Cyborg organoids: Implantation of nanoelectronics via organogenesis for tissue-wide electrophysiology. Nano Lett. 19, 5781–5789 (2019).

13 Lin, Z. et al. Tissue-embedded stretchable nanoelectronics reveal endothelial cell–mediated electrical maturation of human 3D cardiac microtissues. Sci. Adv. 9, eade8513 (2023).

14 Gao, H. et al. Graphene-integrated mesh electronics with converged multifunctionality for tracking multimodal excitation-contraction dynamics in cardiac microtissues. Nat. Commun. 15, 2321 (2024).

15 Tian, B. et al. Macroporous nanowire nanoelectronic scaffolds for synthetic tissues. Nat. Mater. 11, 986–994 (2012).

16 Kalmykov, A. et al. Organ-on-e-chip: Three-dimensional self-rolled biosensor array for electrical interrogations of human electrogenic spheroids. Sci. Adv. 5, eaax0729 (2019).

17 Park, Y. et al. Three-dimensional, multifunctional neural interfaces for cortical spheroids and engineered assembloids. Sci. Adv. 7, eabf9153 (2021).

18 Huang, Q. et al. Shell microelectrode arrays (MEAs) for brain organoids. Sci. Adv. 8, eabq5031 (2022).

19 Choi, S. J. et al. 3D Spatiotemporal Electrophysiology of Cardiac Organoids Using Shell Microelectrode Arrays. Adv. Mater. 38, e06793 (2025).

20 Kim, K. et al. Highly stretchable 3D microelectrode array for noninvasive functional evaluation of cardiac spheroids and midbrain organoids. Adv. Mater. 37, e2412953 (2024).

21 Yang, X. et al. Kirigami electronics for long-term electrophysiological recording of human neural organoids and assembloids. Nat. Biotechnol. 42, 1836–1843 (2024).

22 Fan, J. A. et al. Fractal design concepts for stretchable electronics. Nat. Commun. 5, 3266 (2014).

23 Rubehn, B. & Stieglitz, T. In vitro evaluation of the long-term stability of polyimide as a material for neural implants. Biomaterials 31, 3449–3458 (2010).

24 Engler, A. J. et al. Embryonic cardiomyocytes beat best on a matrix with heart-like elasticity: scar-like rigidity inhibits beating. J. Cell Sci. 121, 3794–3802 (2008).

25 Lind, J. U. et al. Instrumented cardiac microphysiological devices via multimaterial three-dimensional printing. Nat. Mater. 16, 303–308 (2017).

26 Cappello, G. et al. Osmotic pressure induces unexpected relaxation of contractile 3D microtissue. Eur. Phys. J. E 48, 34 (2025).

27 Shilov, S. Y. et al. Biocompatibility of 3D-printed PLA, PEEK and PETG: adhesion of bone marrow and peritoneal lavage cells. Polymers 14, 3958 (2022).

28 Rohr, S. Role of gap junctions in the propagation of the cardiac action potential. Cardiovasc. Res. 62, 309–322 (2004).

29 Spach, M. S., Barr, R. C., Serwer, G. A., Kootsey, J. M. & Johnson, E. A. Extracellular potentials related to intracellular action potentials in the dog purkinje system. Circ. Res. 30, 505–519 (1972).

30 Baer, M., Best, P. M. & Reuter, H. Voltage-dependent action of tetrodotoxin in mammalian cardiac muscle. Nature 263, 344–345 (1976).

31 Herron, T. J. et al. Extracellular matrix-mediated maturation of human pluripotent stem cell-derived cardiac monolayer structure and electrophysiological function. Circ. Arrhythm. Electrophysiol. 9, e003638 (2016).

32 Tertoolen, L. G. J., Braam, S. R., van Meer, B. J., Passier, R. & Mummery, C. L. Interpretation of field potentials measured on a multi electrode array in pharmacological toxicity screening on primary and human pluripotent stem cell-derived cardiomyocytes. Biochem. Biophys. Res. Commun. 497, 1135–1141 (2018).

33 Karbassi, E. et al. Cardiomyocyte maturation: advances in knowledge and implications for regenerative medicine. Nat. Rev. Cardiol. 17, 341–359 (2020).

34 La Manno, G. et al. RNA velocity of single cells. Nature 560, 494–498 (2018).

35 Acun, A., Nguyen, T. D. & Zorlutuna, P. In vitro aged, hiPSC-origin engineered heart tissue models with age-dependent functional deterioration to study myocardial infarction. Acta Biomater. 94, 372–391 (2019).

36 Li, J. et al. Molecular and electrophysiological evaluation of human cardiomyocyte subtypes to facilitate generation of composite cardiac models. J. Tissue Eng. 13, 20417314221127908 (2022).

37 Campostrini, G., Windt, L. M., van Meer, B. J., Bellin, M. & Mummery, C. L. Cardiac tissues from stem cells: new routes to maturation and cardiac regeneration. Circ. Res. 128, 775–801 (2021).

38 Kim, M., Hwang, D. G. & Jang, J. Bioprinting approaches in cardiac tissue engineering to reproduce blood-pumping heart function. iScience 28, 111664 (2025).

39 Feric, N. T. & Radisic, M. Maturing human pluripotent stem cell-derived cardiomyocytes in human engineered cardiac tissues. Adv. Drug Deliv. Rev. 96, 110–134 (2016).

40 Ewoldt, J. K. et al. Induced pluripotent stem cell-derived cardiomyocyte in vitro models: benchmarking progress and ongoing challenges. Nat. Methods 22, 24–40 (2025).

41 Yang, X., Pabon, L. & Murry, C. E. Engineering adolescence: maturation of human pluripotent stem cell-derived cardiomyocytes. Circ. Res. 114, 511–523 (2014).

42 Kang, X., Cheemalamarri, S. K. & Yin, Q. Organoid: a promising solution to current challenges in cancer immunotherapy. *npj Biomed*. Innov. 2, 49 (2025).

43 Gao, H. et al. Bioinspired two-in-one nanotransistor sensor for the simultaneous measurements of electrical and mechanical cellular responses. Sci. Adv. 8, eabn2485 (2022).

44 Kléber, A. G. & Rudy, Y. Basic mechanisms of cardiac impulse propagation and associated arrhythmias. Physiol. Rev. 84, 431–488 (2004).

45 Maltsev, V. A. & Undrovinas, A. Late sodium current in failing heart: friend or foe? Prog. Biophys. Mol. Biol. 96, 421–451 (2008).

46 Bers, D. M. Cardiac excitation–contraction coupling. Nature 415, 198–205 (2002).

47 Fozzard, H. A., Sheets, M. F. & Hanck, D. The sodium channel as a target for local anesthetic drugs. Front. Pharmacol. 2, 68 (2011).

48 Stork, D. et al. State dependent dissociation of HERG channel inhibitors. Br. J. Pharmacol. 151, 1368–1376 (2007).

49 Guzman, J. N. et al. Systemic isradipine treatment diminishes calcium-dependent mitochondrial oxidant stress. J. Clin. Invest. 128, 2266–2280 (2018).

50 Bagley, J. A., Reumann, D., Bian, S., Lévi-Strauss, J. & Knoblich, J. A. Fused cerebral organoids model interactions between brain regions. Nat. Methods 14, 743–751 (2017).

51 Kim, J.-i., et al. Human assembloid model of the ascending neural sensory pathway. Nature 642, 143–153 (2025).

52 Schmidt, C. et al. Multi-chamber cardioids unravel human heart development and cardiac defects. Cell 186, 5587–5605 (2023).

53 Kopinski-Grünwald, O., Guillaume, O., Ferner, T., Schädl, B. & Ovsianikov, A. Scaffolded spheroids as building blocks for bottom-up cartilage tissue engineering show enhanced bioassembly dynamics. Acta Biomater. 174, 163–176 (2024).

54 Huang, C.-J., Schild, L. & Moczydlowski, E. G. Use-dependent block of the voltage-gated Na(+) channel by tetrodotoxin and saxitoxin: effect of pore mutations that change ionic selectivity. J. Gen. Physiol. 140, 435–454 (2012).

55 Viswanathan, P. C., Shaw, R. M. & Rudy, Y. Effects of IKr and IKs heterogeneity on action potential duration and its rate dependence. Circulation 99, 2466–2474 (1999).

56 Burridge, P. W. et al. Chemically defined generation of human cardiomyocytes. Nat. Methods 11, 855–860 (2014).

57 Balafkan, N. et al. A method for differentiating human induced pluripotent stem cells toward functional cardiomyocytes in 96-well microplates. Sci. Rep. 10, 18498 (2020).

