## Supplemental Information for "Flytrap-Inspired Mesh-Trap Bioelectronics for Full Spherical Electrophysiological Interrogation of 3D Tissues"

---

[1] Department of Electrical and Computer Engineering, University of Massachusetts, Amherst, MA, USA.

[2] Department of Polymer Science and Engineering, University of Massachusetts, Amherst, MA, USA.

[3] Institute for Applied Life Sciences, University of Massachusetts, Amherst, MA, USA

[4] Department of Biomedical Engineering, University of Massachusetts, Amherst, MA, USA.

**This file includes:**

Supplementary Figures S1 – S17

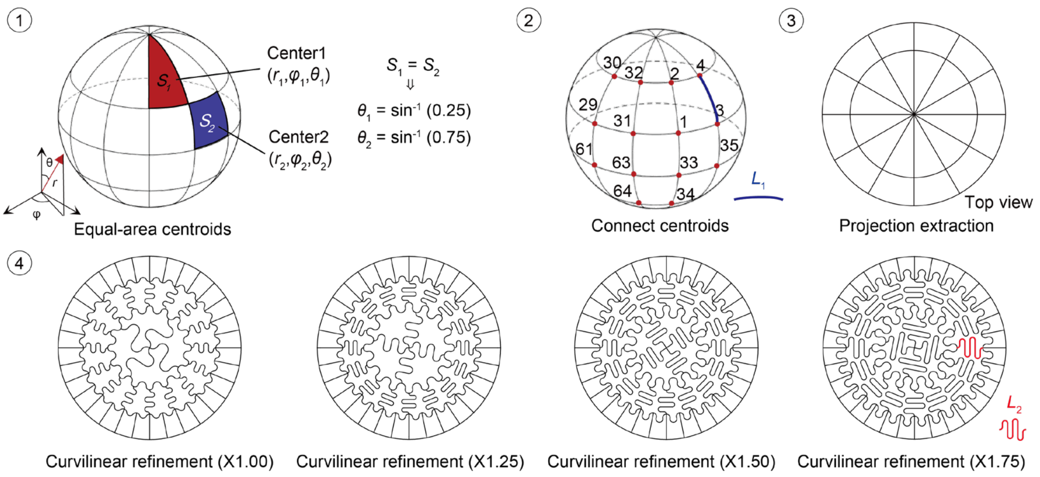


**Fig. S1. Mesh design based on principles of equal-area partitioning and geometric projection.** The spherical surface was subdivided into 64 equal-area regions using latitude-longitude coordinates, and the geometric center of each region was determined (left). Adjacent center points were connected on the sphere, and the corresponding spherical arc length $L_1$ was calculated (middle). The same center points and connections were projected onto the XY plane to obtain the planar distance $L_2$. Each planar segment was then replaced by an S-shaped curve whose contour length matched $L_1$ (correction factor = 1) (right). To account for finite mesh linewidth, additional correction factors of 1.25, 1.5, and 1.75 were applied to elongate the S-shaped curves, and 1.75 was selected based on mechanical simulations (bottom). Detailed procedures are provided in the *Methods*.

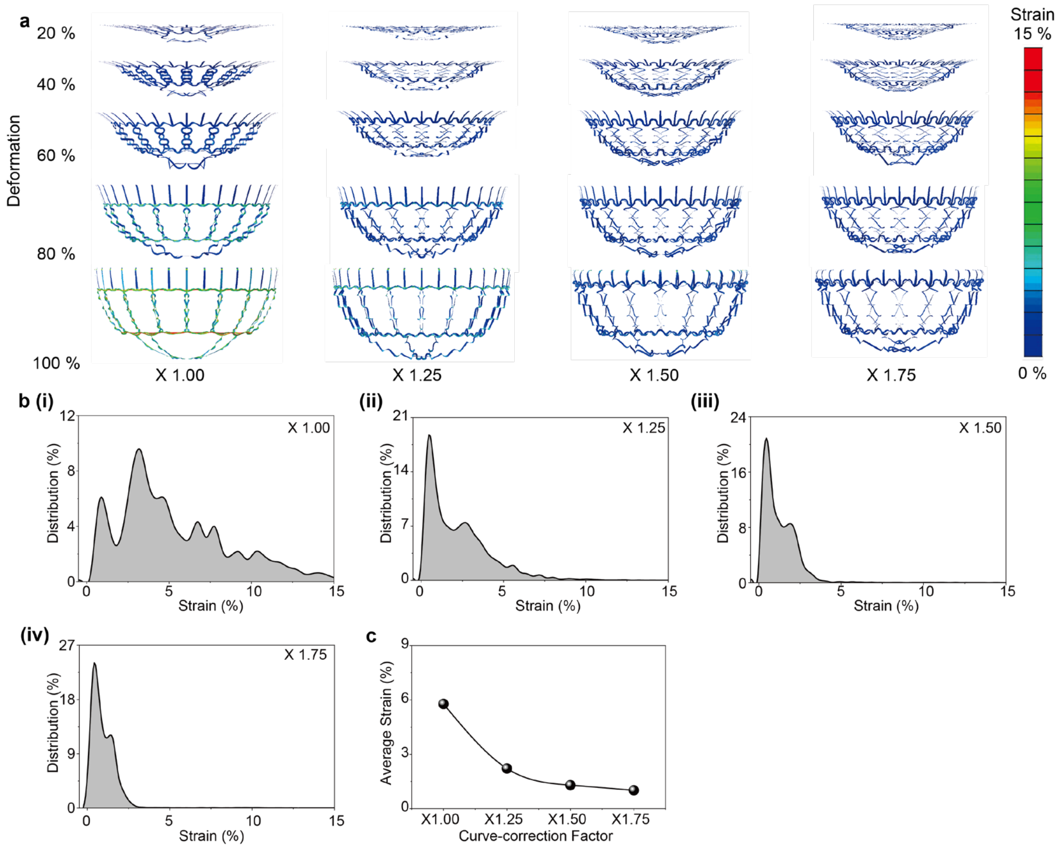


**Fig. S2. Mesh structural optimization by Finite element analysis. a**, Strain distributions under different deformation conditions for varying curve-correction factors. For different curve-correction factors, the mesh gradually conforms more closely to the spherical contour. However, when the factor is 1.00, the mesh exhibits pronounced distortion, which supports the necessity of introducing curve-correction factors to optimize the design. **b**, Probability density distributions of local strain for different curve-correction factors **(i)** 1.00, **(ii)** 1.25, **(iii)** 1.50, and **(iv)** 1.75. **c**, Average strain as a function of the curve-correction factor.

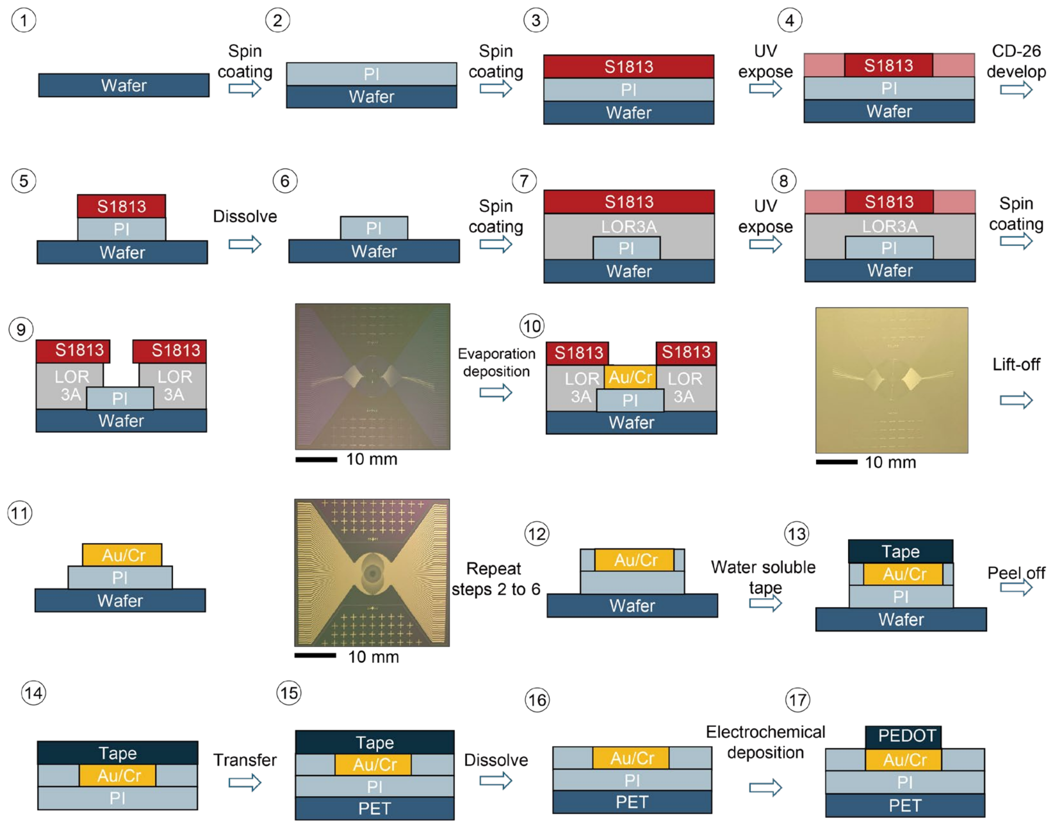


**Fig. S3. Mesh device fabrication.** Device fabrication was performed using standard microfabrication techniques. Silicon wafers with a 300-nm $SiO_2$ layer were used as substrates, followed by spin-coating and curing of the polyimide (PI) structural layer. Photolithography with S1813 photoresist was used to define the PI geometry, including UV exposure, CD-26 development, and solvent cleaning. A bilayer lift-off process (LOR 3A/S1813) was then applied to pattern the electrode region. Cr/Au (10 nm/100 nm) was deposited by e-beam evaporation and defined by lift-off. Additional PI coating and photolithography steps were repeated to form a mesh architecture with the metal layer embedded between top and bottom PI passivation layers. After HF etching of the $SiO_2$ sacrificial layer, the device was released using water-soluble tape and transferred onto a PET substrate, followed by tape dissolution in water. Finally, PEDOT:PSS was electrodeposited onto the electrodes to improve electrochemical performance. Detailed fabrication parameters are provided in the *Methods*.

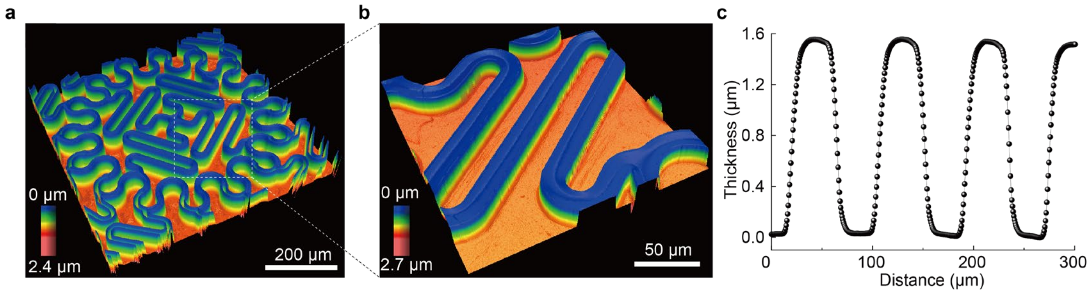


**Fig. S4. Device surface topography. a**, 3D surface topography of the mesh structure. **b**, Magnified three-dimensional surface topography. **c**, Surface profiles extracted along the indicated directions. **(c)**, 2D surface profilometer revealed an average device thickness of approximately 1.6 μm.

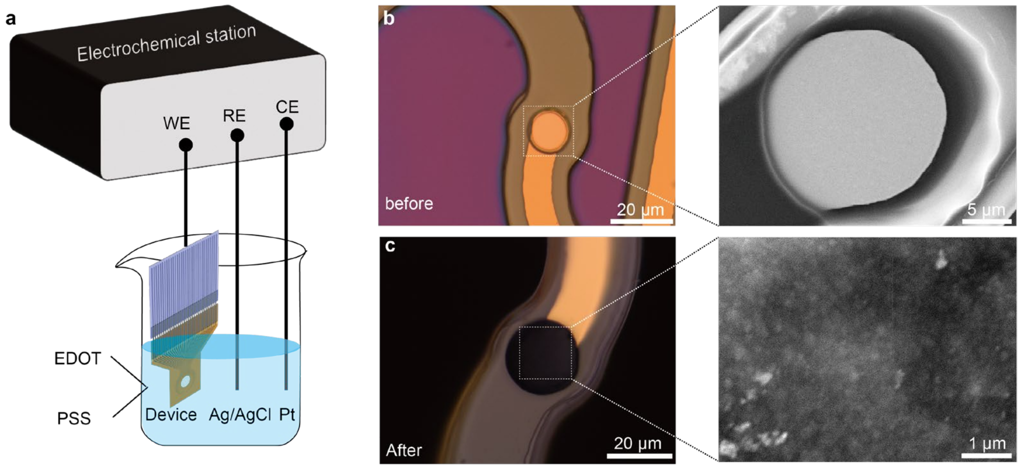


**Fig. S5. Electrochemical deposition of PEDOT:PSS. a**, Schematic illustration of the electrochemical deposition setup, showing the three-electrode configuration (working electrode (WE), reference electrode (RE), and counter electrode (CE)). **b**, Optical image of the electrode site before deposition. **c**, Optical image of the same electrode site after deposition. Right panels show scanning electron microscopy (SEM) images of the electrode surface before and after deposition. The electrode surface is smooth and flat before deposition, whereas after deposition it exhibits a rough, high surface area morphology.

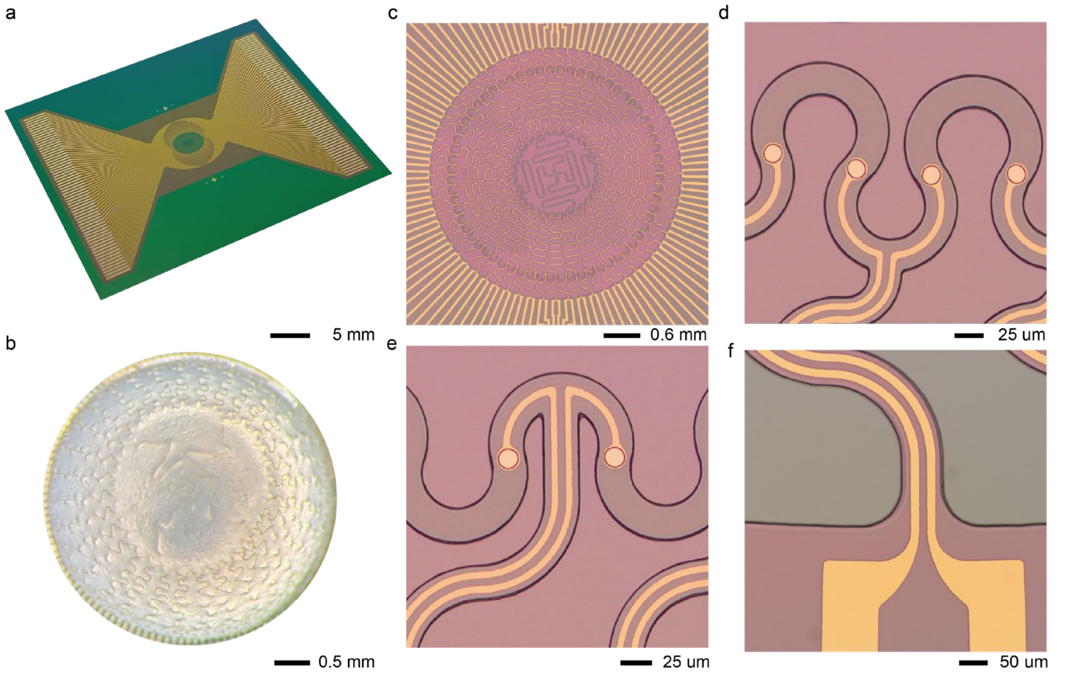


**Fig. S6. Mesh trap integrated with 256 electrodes. a**, Perspective view of the fabricated device with an overall dimension of 34 × 37 mm. **b**, Top view of the device on a plastic sphere. (**c–f**), Magnified views of device regions. For improved electrode density, two interconnects are integrated on an individual ribbon. This strategy ensures higher integration density without increasing the overall device size. (c) is a stitched image due to the large device size.

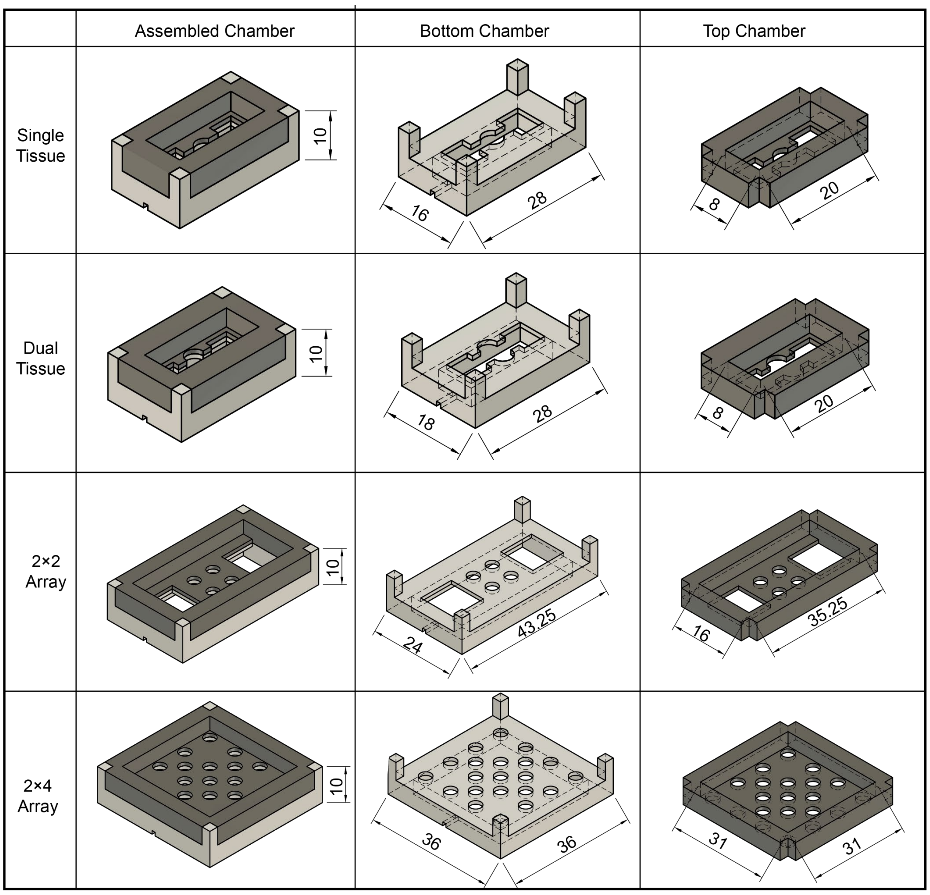


**Fig. S7. Schematic illustration of the chamber design used in this study.** (From top to bottom) The chambers designed for a single-mesh trap (10 × 16 × 28 mm, with an estimated volume of approximately 1.5 mL), a dual-mesh (10 × 18 × 28 mm, with an estimated volume of approximately 1.5 mL), 2×2 mesh trap array (10 × 24 × 43 mm, with an estimated volume of approximately 5 mL), and 2×4 mesh trap array (10 × 36 × 36 mm, with an estimated volume of approximately 9 mL), respectively.

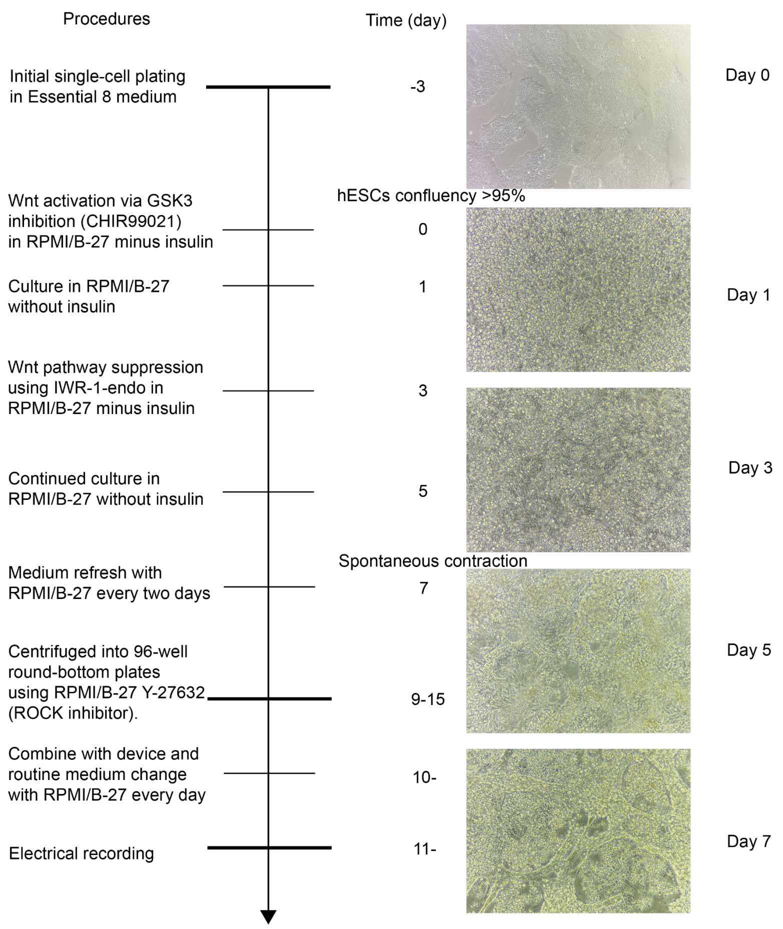


**Fig. S8. Timeline for microtissue culture.** Representative bright-field microscopy images of cardiac microtissues recorded at different culture stages. The tissue surface gradually transitioned from a relatively smooth and compact morphology to a rougher, heterogeneous structure with increased texture and visible cellular remodeling, indicating progressive structural reorganization during long-term culture. Detailed culture conditions and imaging parameters are provided in the *Methods*.

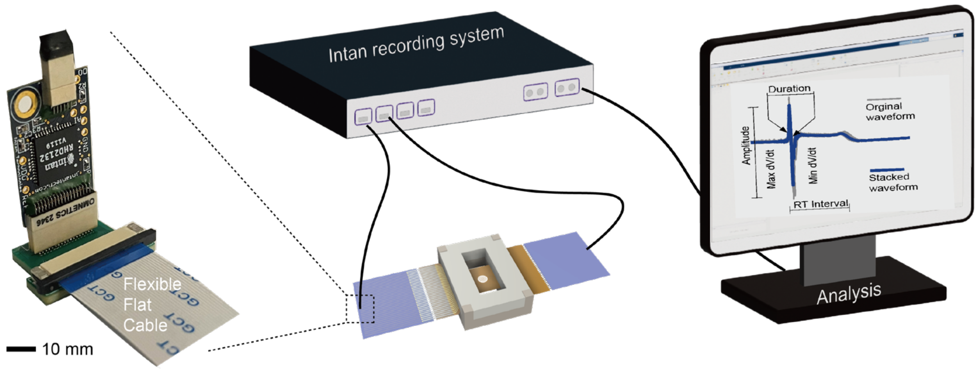


**Fig. S9. Schematic overview of the electrophysiological recording.** Electrical recordings were performed using an Intan RHD system with manufacturer-provided cables and acquisition software. The headstage was interfaced with the device through a custom PCB incorporating a 32-channel flexible flat cable converter. Signals were acquired at 20 kHz. All measurements were conducted inside a custom electromagnetic shielding enclosure on a vibration-isolated optical table to minimize noise. Data analysis was performed in MATLAB R2024a using both Intan file conversion tools and custom scripts. Detailed procedures are provided in the *Methods*.

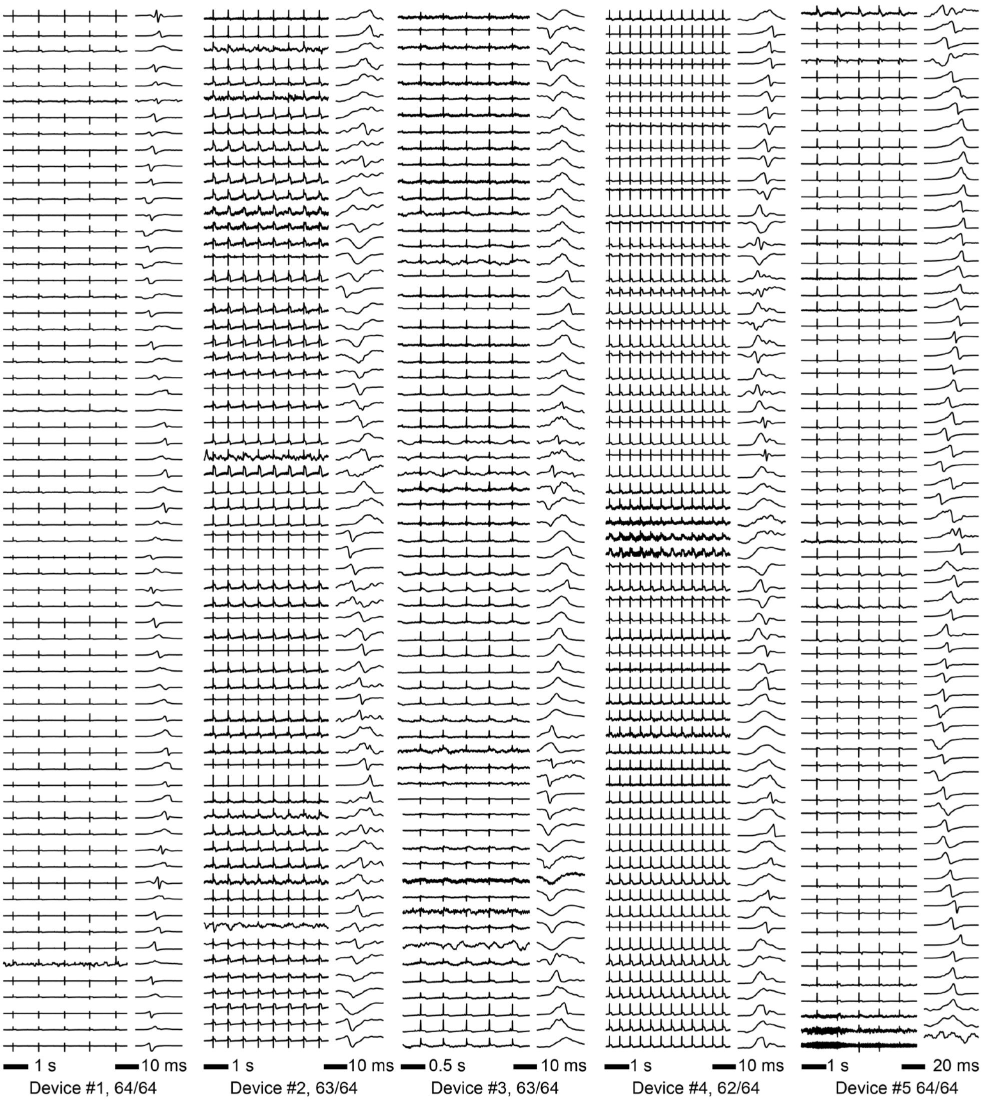


**Fig. S10. Representative data of five different devices.** Representative recordings from five mesh-trap devices, each integrated with 64 recording electrodes. The recordings showed an overall yield >97%. Clear inter-channel time delays were consistently observed, providing the basis for subsequent spatial propagation and conduction analysis.

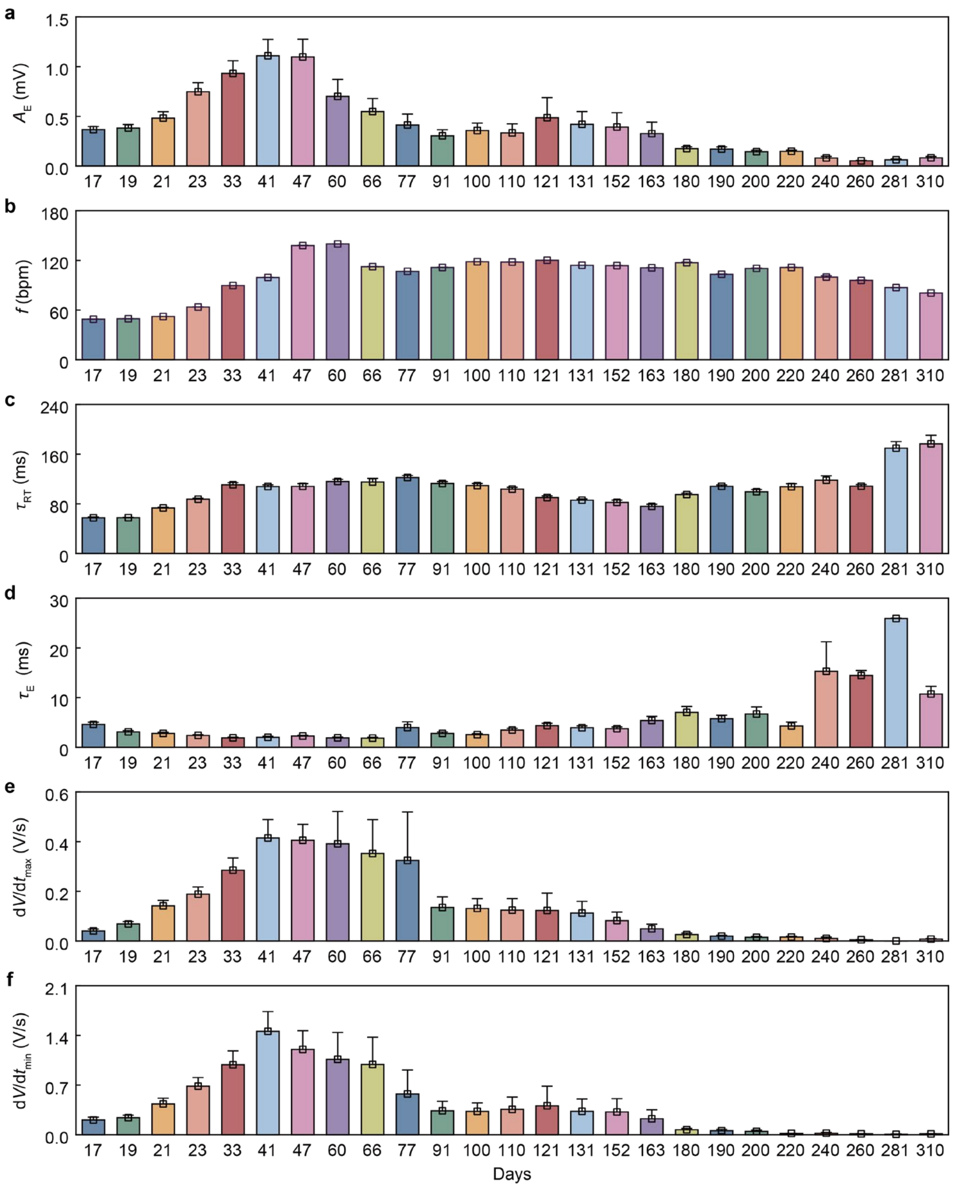


**Fig. S11. Evolution of electrophysiological metrics over 300 days. a**, Beating frequency ($f$). **b**, Signal amplitude ($A_E$). **c**, RT interval ($\tau_{RT}$). **d**, Duration ($\tau_E$). **e**, Maximum derivatives of the voltage signals ($dV/dt_{max}$). **f**, Minimum derivatives of the voltage signals ($dV/dt_{min}$). Most features exhibited a gradual decline after day 47, consistent with the typical aging-like time described in the main text.

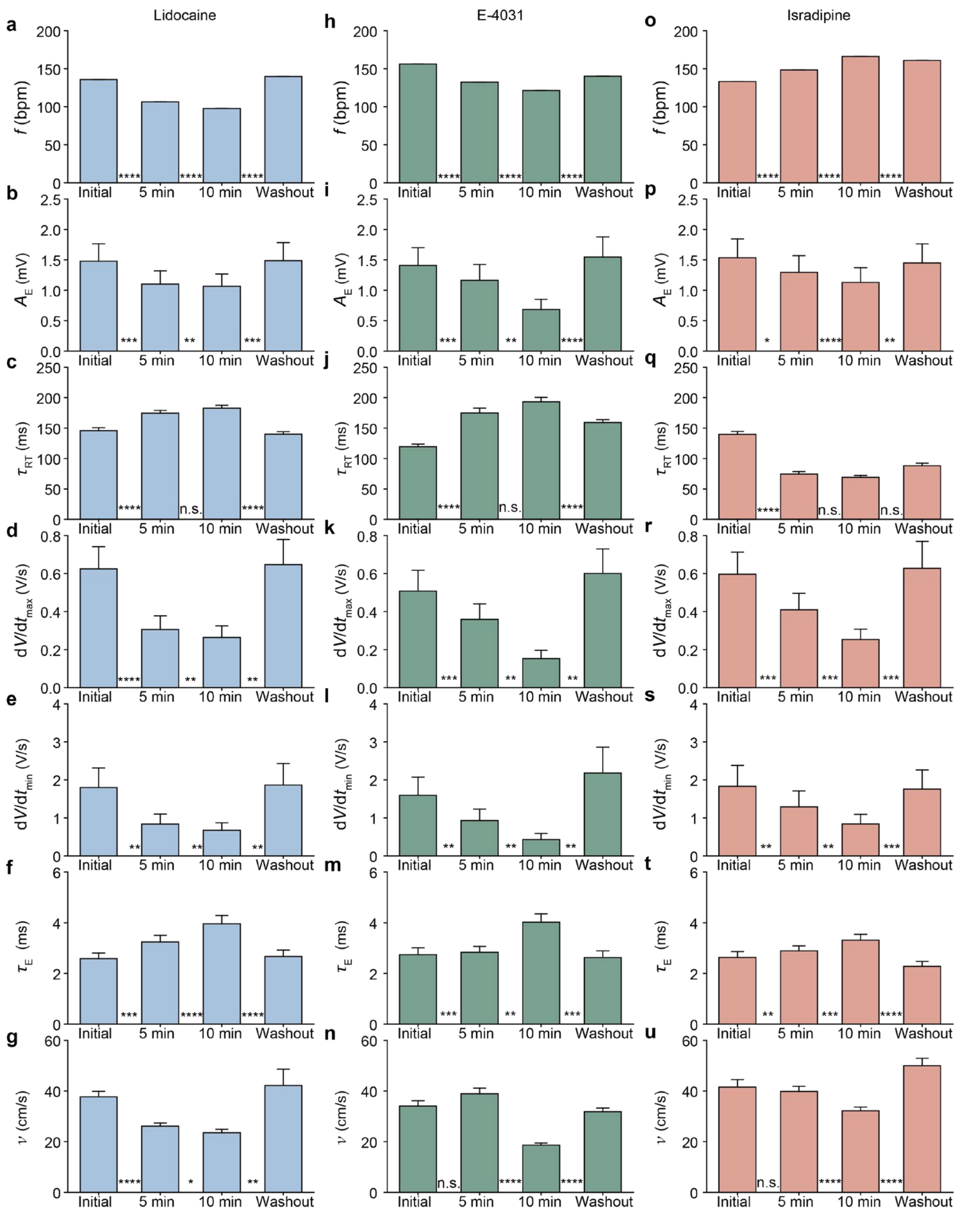


**Fig. S12. Pharmacological modulation of ion-channel activities.** Evolution of beating frequency ($f$), signal amplitude ($A_E$), RT interval ($\tau_{RT}$), maximum derivatives of the voltage signals ($dV/dt_{max}$), minimum derivatives of the voltage signals ($dV/dt_{min}$), duration ($\tau_E$) and conduction velocity ($v$). **a–g**, Lidocaine ($Na^+$ block, 20 µM). **h–n**, E-4031 ($K^+$ block, 50 nM). **o–u**, Isradipine ($Ca^{2+}$ block, 20 nM). The data are presented as mean ± s.e.m for all, using two-tailed paired t test.

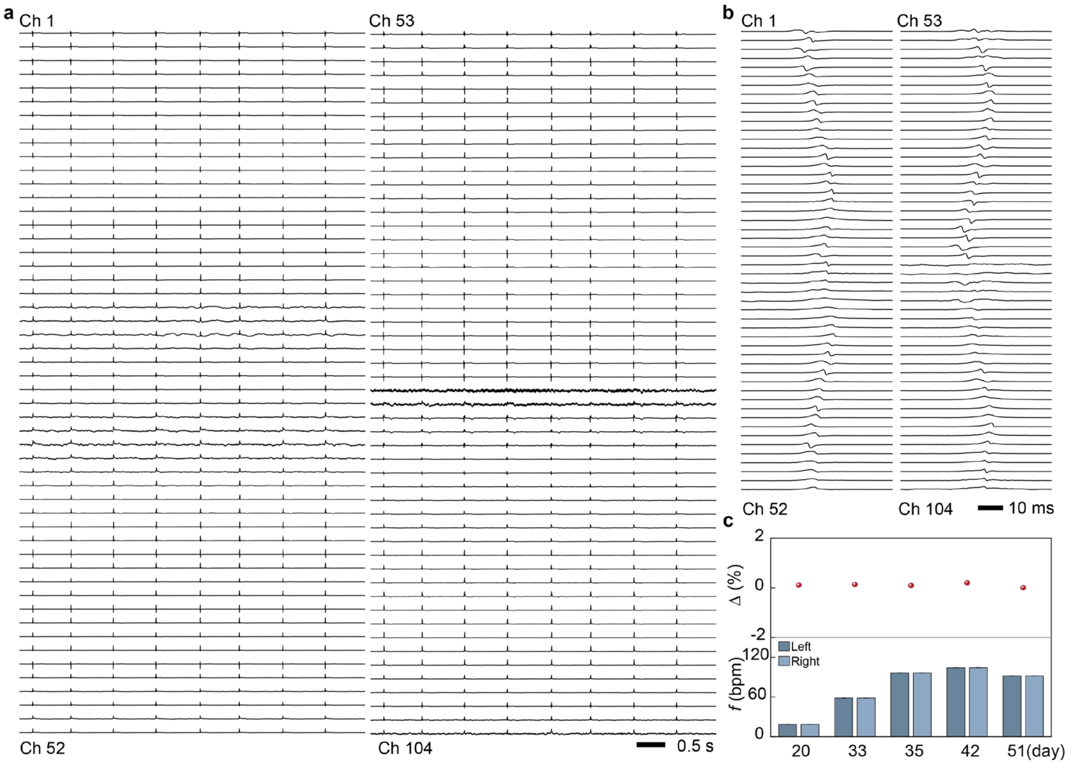


**Fig. S13. Electrophysiological recordings in an assembloid. a**, Representative recordings showing periodic signals across 104 channels. **b**, Locally magnified electrophysiological waveforms. **c**, Comparisions of the temporal evolution of beating frequency (*f*) between the left and right microtissues. The data are presented as mean ± s.e.m. for all.

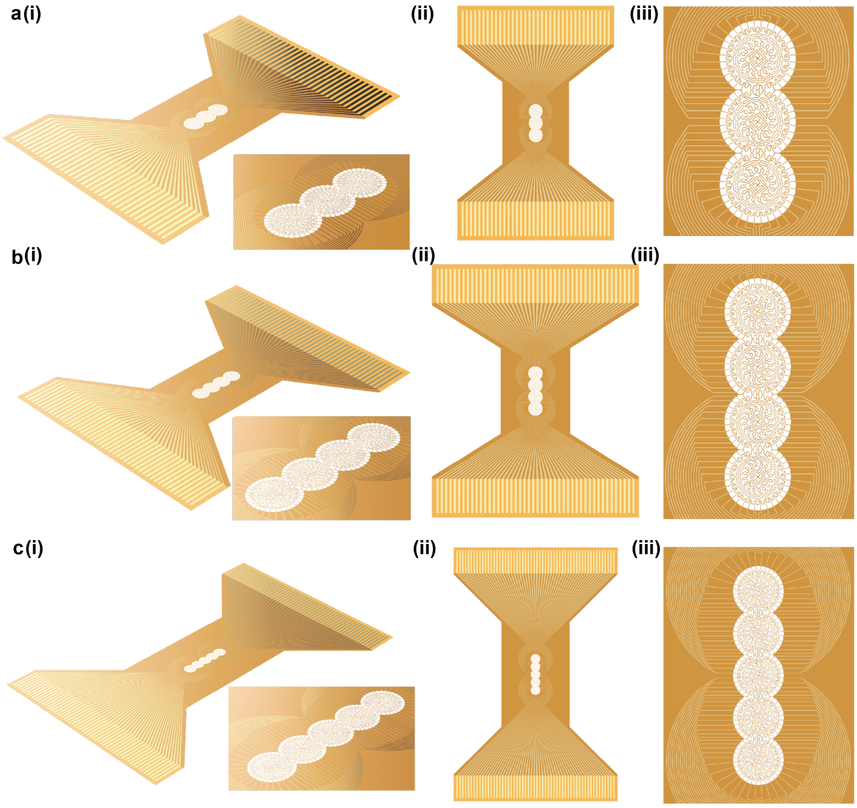


**Fig. S14. Mesh designs for assembloids from three, four, and five tissues. a**, Design for three tissues with a total of 144 electrodes on the top and bottom layers (27 mm × 19 mm). **b**, Design for four tissues with a total of 184 electrodes on the top and bottom layers (30 mm × 24 mm). **c**, Design for five tissues with a total of 224 electrodes on the top and bottom layers (45 mm × 29 mm). **(i)** Three-dimensional view. **(ii)**, Top view. **(iii)**, Magnified view.

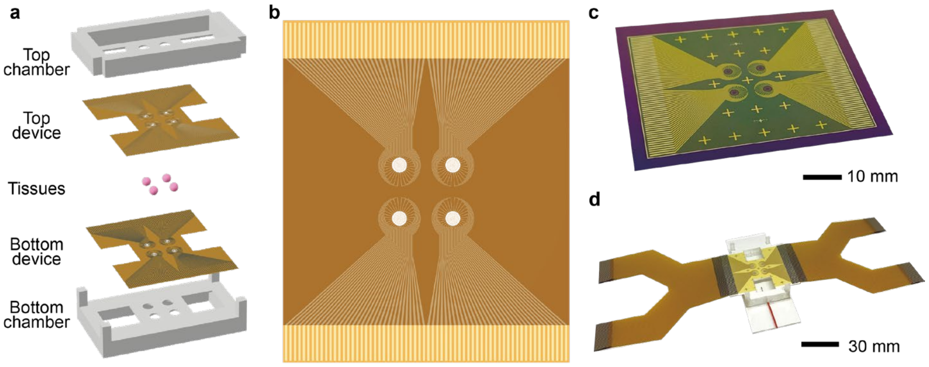


**Fig. S15. 2×2 mesh-trap array. a**, Schematic illustration of the chamber assembly. **b**, Top-view schematic of the 2×2 array with a total of 256 electrodes on the top and bottom layers (39 mm × 32 mm). **c**, Photograph of the fabricated 2×2 array device. **d**, Photograph of the assembled 2×2 array integrated with the chamber.

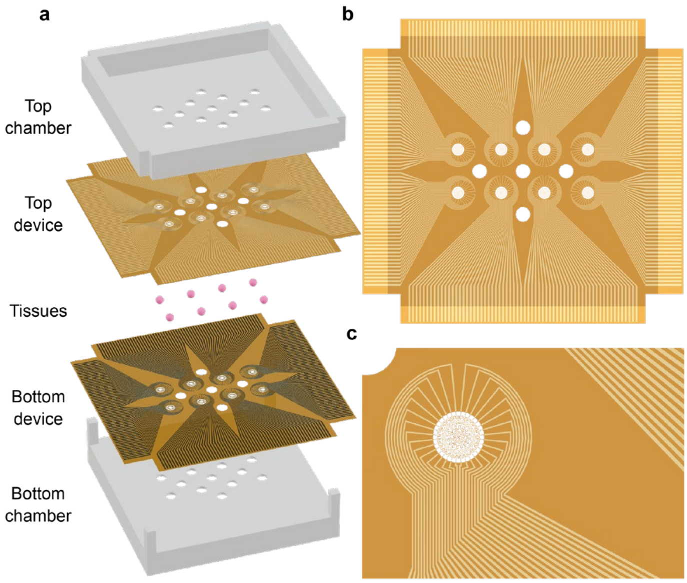


**Fig. S16. 2×4 mesh trap array. a**, Schematic illustration of the chamber assembly with dimensions of 10 × 37 × 37 mm and an estimated volume of approximately 9 mL. **b**, Top-view schematic of the 2×4 array with a total of 512 electrodes on the top and bottom layers (45 mm × 42 mm). **c**, Magnified schematic of the 2×4 array.

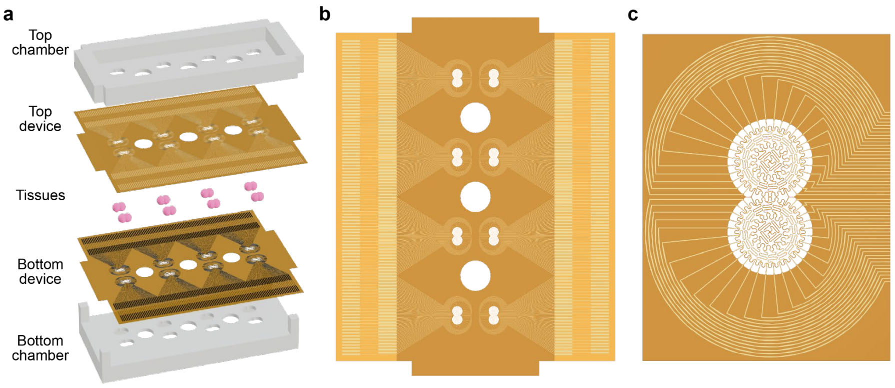


**Fig. S17. 2×4 dual-mesh trap array. a**, Schematic illustration of the chamber assembly with dimensions of 10 × 26 × 59 mm and an estimated volume of approximately 8 mL. **b**, Top-view schematic of the 2×4 array for dual-mesh configuration with a total of 832 electrodes on the top and bottom layers (59 mm × 46 mm). **c**, Magnified schematic of the 2×4 dual-mesh array with a total of 104 electrodes on the top and bottom layers.
